# Superior colliculus visual neural sensitivity at the lower limit of natural self-induced image displacements

**DOI:** 10.1101/2022.06.26.497631

**Authors:** Ziad M. Hafed, Chih-Yang Chen, Fatemeh Khademi

## Abstract

Visual pattern analysis relies on computations from neurons possessing spatially confined receptive fields. Often, such receptive fields are orders of magnitude larger than the visual pattern components being processed, as well as these components’ minute displacements on the retina, whether due to small object or self motions. Yet, perception effortlessly handles such visual conditions. Here, we show that in the primate superior colliculus, a brain structure long associated with oculomotor control, neurons with relatively large receptive fields are still sensitive to visual pattern displacements as small as 1 min arc. We used real-time gaze-contingent retinal image stabilization to control the instantaneous spatio-temporal luminance modulation of detailed patterns experienced by neurons, probing sensitivity to the lower limit of natural self-induced image displacements. Despite a large difference between pattern displacement amplitudes and receptive field sizes, collicular neurons were strongly sensitive to the visual pattern consequences of the smallest possible image-shifting eye movements.

## Introduction

Image analysis in the primate visual system is performed by neurons having individual receptive fields (RF’s) sampling confined regions of the retinal image. Outside the fovea, and particularly in higher visual areas, RF’s can be large. This is also the case in sensory-motor structures like the superior colliculus (SC) ^1-3^, which itself has a rich and diverse visual repertoire ^1-7^.

Integration of relatively large image regions by individual RF’s raises questions about how detailed visual pattern analysis can occur when the local features inside an RF are much smaller than RF size. Among these questions is what nature of visual processing takes place in the SC, a sensory-motor structure, when compared to other visual areas that are more distant from the motor control apparatuses. For example, the SC contributes to saccade generation ^8^, and its diverse visual properties seem to be optimized for detecting stimuli for the purpose of gaze orienting ^9^. Does this mean that SC neurons are incapable of detailed visual pattern analysis that may be more the purview of visual cortex?

We investigated this question by exploring whether SC neurons are sensitive to the visual pattern consequences of minute image displacements over their RF’s. With the head fixed, a lower limit on natural self-induced retinal image motion is that caused by slow ocular position drifts during gaze fixation (Fig. 1A) ^10-12^. With stable external stimuli, such drifts introduce image pattern displacements over individual RF’s that are much smaller than the RF’s themselves (Fig. 1B). Thus, the local pattern features of the stimuli never really leave the RF’s during drifts. Yet, theoretical and perceptual work suggests that small displacements associated with ocular position drifts reformat images in meaningful ways for perception ^13-16^. The reformatting itself is a direct consequence of eyeball rotation: it is a property of the image formation process. However, for the reformatting to be effective for perception, downstream neural processing stages need to be also sensitive to them. Therefore, we asked whether SC neurons functionally utilize the visual reformatting afforded by ocular position drifts in their activity (Fig. 1B).

**Figure 1.**
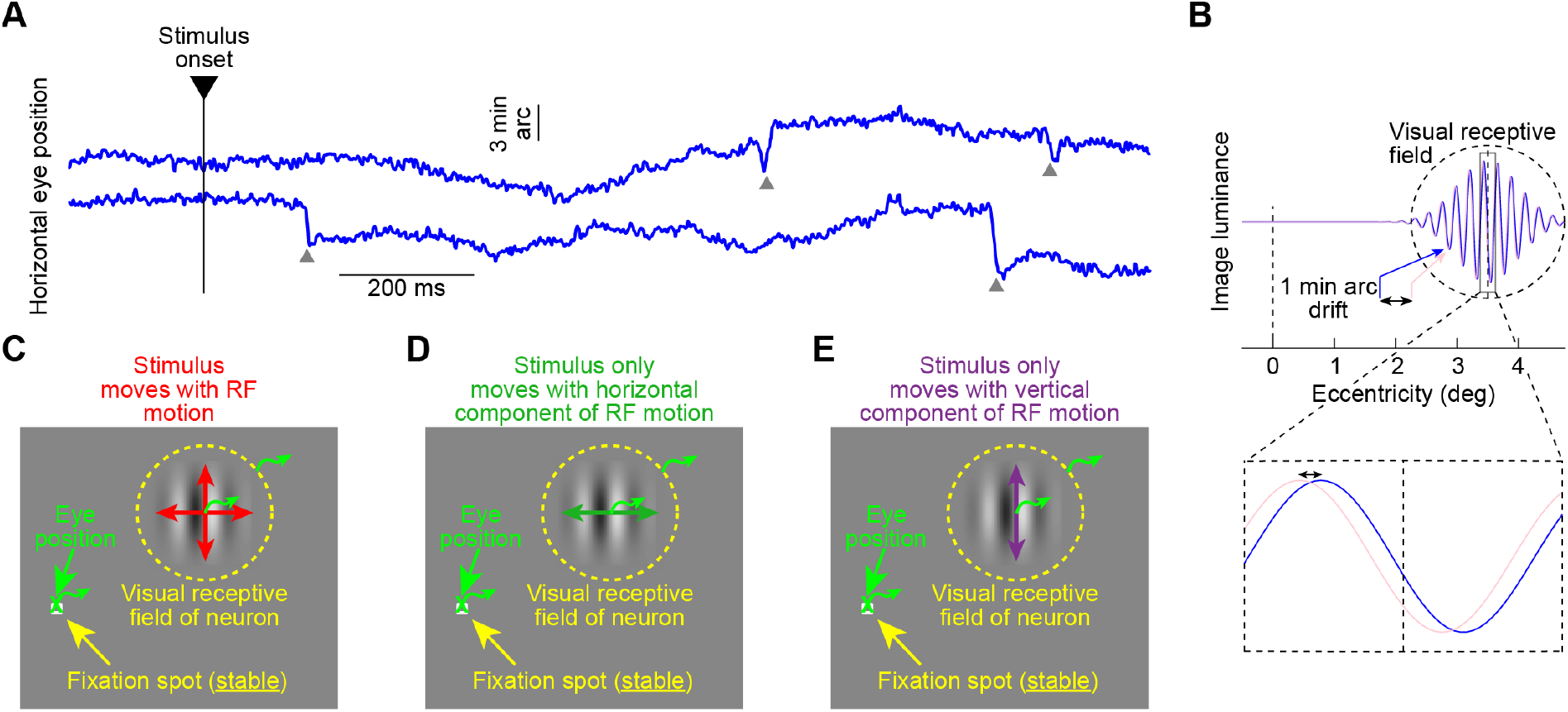
Isolating the visual-pattern consequences of slow eye position drifts on SC neural activity. **(A)** Horizontal eye position from two example gaze fixation trials. Microsaccades are highlighted with arrows and were removed from all analyses (Methods). In between, slow eye position drifts occurred with a retinal-image displacement range on the order of 1-3 min arc. **(B)** The luminance profile of a 4.44 cpd gabor grating placed within a visual receptive field (RF) of a hypothetical SC neuron for two instantaneous eye positions separated by 1 min arc (blue versus pink; the bottom inset magnifies one cycle). The image displacement is much smaller than the RF size (depicted based on our prior measurements ^3,9^). **(C)** To characterize SC neural sensitivity to pattern displacements on the order of 1-3 min arc, we exploited the slow position drifts in **A**. Monkeys fixated a stable spot, and we continuously translated the image of an eccentric grating with fixational eye motion, thus stabilizing the grating’s image within the continuously moving retinotopic RF. This minimized the displacements of the grating in the RF that would have otherwise occurred. **(D)** In other conditions, we only stabilized the horizontal component of ocular position changes (minimizing RF image shifts orthogonal to the grating). **(E)** In yet other conditions, we only stabilized the vertical component of eye drift. Curly bright green arrows in **C, D, E** schematize the eye position changes and their associated retinotopic RF motions.

We employed the technique of gaze-contingent retinal image stabilization, combined with grating images of different properties, to identify a direct SC neural correlate of perceptual effects ^13^ associated with ocular position drifts. We presented gratings to the RF’s of SC neurons either stably on the display (and thus moving on the retina due to drifts) or using gaze-contingent display updates, as was done earlier in the primary visual cortex with simple spot and bar stimuli ^17^. We used, instead, patterned gratings and different experimental manipulations of retinal image motions to demonstrate clear SC neural sensitivity (even extrafoveally) to the diminutive visual pattern consequences of ocular position drifts. These results complement studies in the retina highlighting the impact of visual reformatting by drifts on the retinal output ^18^, and they demonstrate that SC neurons can contribute to visual scene analysis with high fidelity despite their relatively large RF’s.

## Results

We asked whether visually-responsive SC neurons are sensitive to image pattern displacements at the lower limit of natural self-induced motion. We did so by recording SC neural activity from head-fixed monkeys precisely holding their gaze on a small, stable fixation spot of 8.5 × 8.5 min arc dimensions. During fixation, the eyes drifted slowly ^10-12^, causing retinal image displacements on the order of 1 min arc magnitude (Fig. 1A), with occasional larger microsaccades. The scale of retinal image displacements associated with ocular position drifts, as small as the approximate distance between two individual foveal cone photoreceptors, is considerably smaller than SC RF sizes, especially extrafoveally ^1-3,9^. It is also frequently much smaller than the viewed image patterns themselves. Consider, for example, a gabor grating of 4.44 cycles/deg (cpd) in an RF of a neuron preferring 3.5 deg eccentricity. A displacement over the RF of the grating’s retinal image by 1 min arc would cause minimal change to the overall luminance pattern experienced by the neuron (Fig. 1B). Yet, theoretical considerations suggest that such minute pattern displacements can still matter for perception ^10,13-16^. We, therefore, investigated whether SC neurons are sensitive to these diminutive pattern displacements.

We utilized real-time retinal image stabilization to move a visual pattern (gabor grating) in lock with instantaneous eye position (Methods). We compared neural activity with a stable pattern on the display (thus moving with respect to a continuously moving retinotopic RF) to neural activity with a moving pattern on the display tracking the eye movements (and thus rendered more stable with respect to the RF; Fig. 1C). If the SC is sensitive to visual pattern consequences of retinal image displacements as small as those in Fig. 1A, B, then the neural responses should differ between the two conditions. Critically, the fixation spot was always stable on the display, allowing the monkeys to properly anchor their gaze independently of retinal image stabilization. Indeed, the characteristics of both ocular position drifts and microsaccades across all of our gaze-contingent manipulations (Methods) were unaltered by whether the grating was stable on the display (control) or not (Figs. S1-S3). This is consistent with evidence that, in steady-state fixation, microsaccades and ocular position drifts act to optimize eye position at the fixated target ^19-22^. Therefore, we experimentally controlled the subtle image displacements of visual patterns over RF’s (Fig. 1), but without altering the natural gaze behavior itself.

### Superior colliculus neurons are sensitive to visual pattern displacements on the order of 1 min arc

We first established the effectiveness of our manipulation. After identifying a visually-responsive neuron, we estimated its retinotopic RF hotspot location and extent (Methods). If we then pegged the stimulus, via retinal image stabilization, at the estimated hotspot location, then the neuron would consistently experience an optimal stimulus. This is in contrast to control trials, during which fixational eye movements could, at any one moment in time, displace the stimulus from the optimal RF hotspot location or otherwise blur it. Thus, neural activity was expected to be elevated with retinal image stabilization (Fig. 2). Alternatively, if we pegged the stimulus at a sub-optimal location relative to the RF during retinal image stabilization, then the neuron’s activity was expected to decrease, because in control trials, eye movements could momentarily bring the stimulus to a more optimal RF position (Fig. S4). These effects are similar to those observed in V1 with the retinal image stabilization technique and simple spot and bar stimuli ^17,23,24^, and they meant that we were now in a good position to explore, in more detail, the visual pattern consequences of minute ocular position drifts on SC image representations.

**Figure 2.**
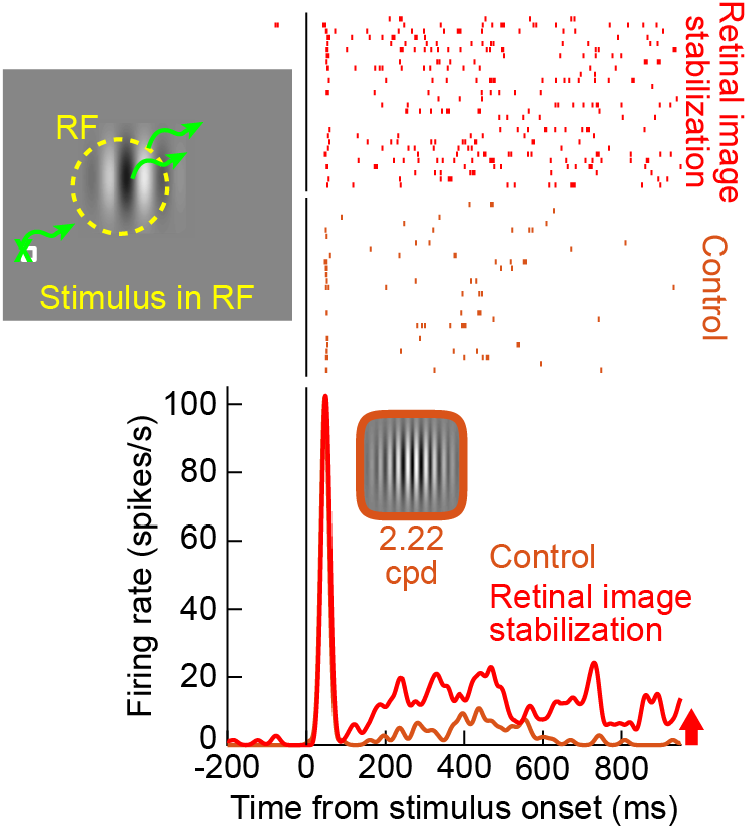
Example SC neuron illustrating the effect of retinal image stabilization on SC activity. In control trials, we presented a grating of middle spatial frequency (2.22 cpd) near RF center. The neuron had a weak response after the initial visual burst. In retinal image stabilization trials, we continuously updated the position of the grating with fixational eye position (predominantly ocular drift; Fig. 1A), to render the grating’s retinotopic position relative to the RF more stable than in control trials. The neuron’s sustained visual response was significantly elevated, suggesting that smearing of the grating position over the RF in control trials reduced the overall responsiveness of the neuron. Our analyses focused on sustained responses, to avoid the transients associated with stimulus onsets at the beginnings of trials (Methods). Fig. S4 also shows an additional validation of the stabilization technique, this time when placing the grating away from the RF center of a recorded neuron.

In all of our subsequent analyses, we only focused on situations like in Fig. 2, with an optimally placed RF stimulus, and also with primarily extrafoveal neurons with RF’s larger than the scale of ocular position drifts; this was the relevant scenario for the questions raised by Fig. 1B. We also excluded all epochs around microsaccades (Methods), because retinal image stabilization with discretized display update times (Methods) is expectedly ^17^ least effective for these faster eye movements (but see Fig. S5 for evidence of the effectiveness of the technique in tracking eye motions even with microsaccades).

Retinal image stabilization of a stimulus at the RF hotspot location consistently elevated SC neural activity, suggesting sensitivity to local image pattern statistics within the RF’s (like those schematized in Fig. 1B). In Fig. 3A, we plotted the normalized population activity after the onset of a 4.44 cpd grating in the RF’s, either in control or with full retinal image stabilization. We found a persistently elevated sustained response (after an initial visual burst due to stimulus onset) for as long as the stimulus was stabilized over the RF’s (compare control to retinal image stabilization firing rates). Given the spatial scale of our image displacements associated with ocular position drifts and the predicted luminance modulations that they introduced (Figs. 1A, S1, S3C-F), this implies that SC neurons can indeed detect minute image pattern displacements much smaller than RF sizes, and also smaller than the pattern features themselves (Fig. 1B). Figure 3B also shows the distribution of individual neural modulation indices (Methods) across our population, with a significant positive shift indicating consistently elevated firing rates during retinal image stabilization (21.74% average modulation index relative to control; p=3.2073×10^−8^; 1-sample t-test; n=61 neurons). Moreover, Fig. S6A shows the individual neuron raw firing rates during sustained fixation (shaded gray region on the x-axis of Fig. 3A, and excluding microsaccades) in the two conditions: practically all neurons exhibited elevated activity for vertical 4.44 cpd gratings stabilized over their RF’s as opposed to being jittered by ocular position drifts in the control condition. This is a direct SC neural correlate of the concept of temporal encoding by ocular position drifts predicted theoretically ^13-15^, but now being viewed from the individual neuron perspective.

**Figure 3.**
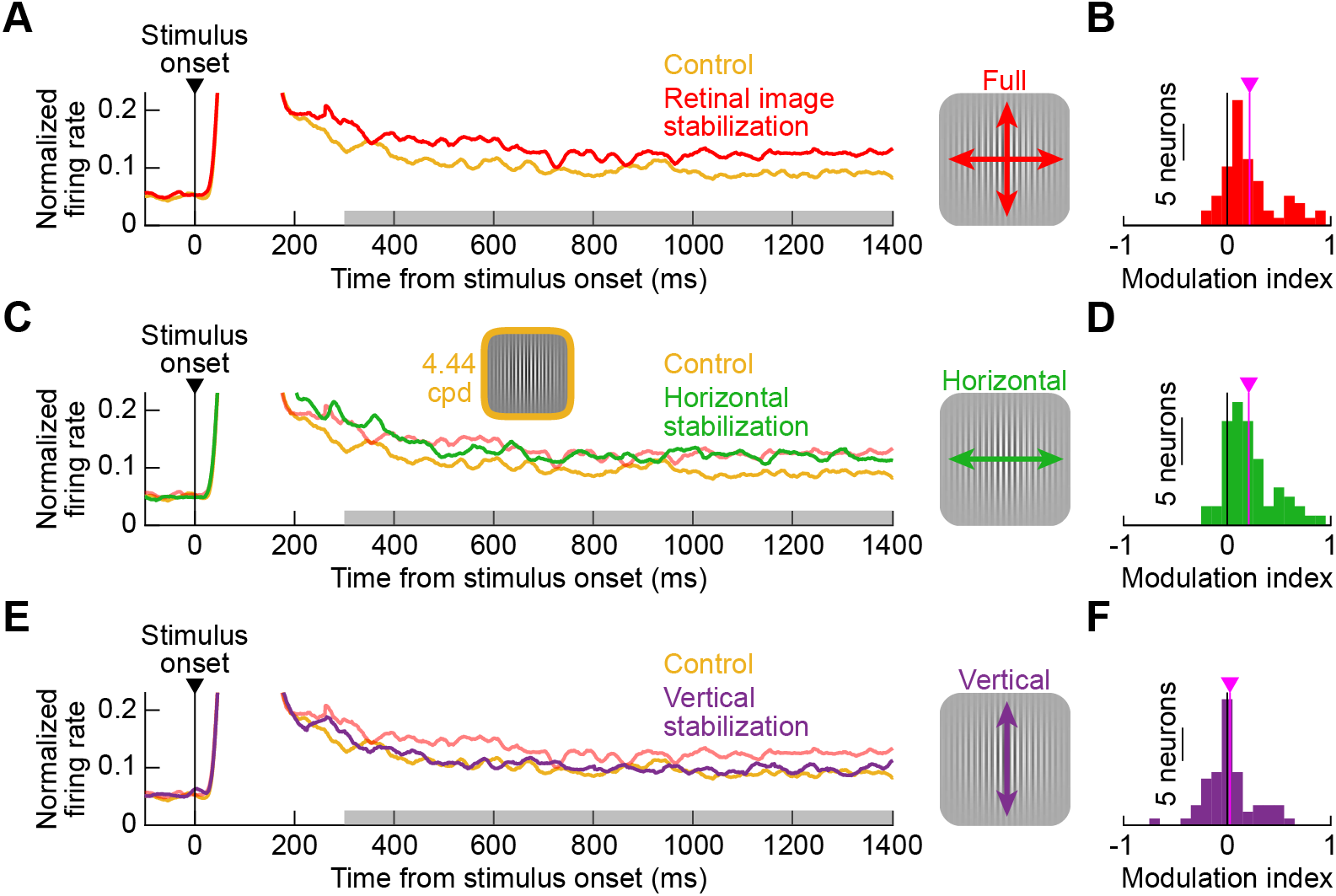
SC neurons are sensitive to image pattern displacements as small as 1-3 min arc in amplitude. **(A)** Normalized firing rate across our population after the onset of a 4.44 cpd grating in the RF’s. Yellow shows the control condition in which the grating was stable on the display, and therefore continuously being displaced over the retinotopic RF’s. The neurons exhibited a sustained response above baseline, as long as there was a stimulus presented. Red shows the full retinal image stabilization condition (Fig. 1C), in which the firing rate was persistently elevated. The gray horizontal bar on the x-axis defines our measurement interval for our analyses (Methods). **(B)** Distribution of per-neuron modulation indices comparing the sustained response under full retinal image stabilization to the sustained response in control (Methods). The vertical pink line shows the mean modulation index across the population (individual neuron raw measurements are also shown in Fig. S6A). **(C)** Same as **A** but for retinal image stabilization of only the horizontal component of eye position displacements (orthogonal to the grating orientation). The neurons were as affected as in the full retinal image stabilization condition (the faint red curve is replicated from **A** for easier comparison). This is because a vertical ocular position drift over a vertical grating causes no luminance modulations over the RF’s beyond the original image pattern. **(D)** Modulation index distribution for the data in **C**, showing similar effects to full retinal image stabilization. **(E)** With vertical retinal image stabilization, the neurons were unaffected by the gaze-contingent image manipulation: the horizontal component of RF motion relative to the grating was the same as in control, resulting in the same neural response. **(F)** Modulation indices for vertical retinal image stabilization were not significantly different from zero across the population.

### Drift-scale pattern displacements orthogonal to the patterns’ orientations drive neural modulations the most

The results of Fig. 3A, B might have been an artifact of world-centered grating motion, which was necessarily introduced by retinal image stabilization (by stabilizing the retinal image, we had to move the stimulus on the display). We, therefore, tested the stronger predictions afforded by either horizontal or vertical retinal image stabilization (Fig. 1D, E). Here, with the same vertical 4.44 cpd grating, if we stabilized only the horizontal component of fixational eye movements, then any residual vertical movements of the grating over the retinotopic RF’s (uncompensated by the partial gaze-contingent technique) would not alter the luminance pattern over the RF’s too much; this is because the vertical component of ocular position drifts is much smaller than the grating size and also parallel to it (Fig. 1D). On the other hand, if we stabilized only the vertical component of fixational eye movements (Fig. 1E), then the residual (uncompensated) horizontal movements of the grating over the retinotopic RF’s would cause luminance pattern changes like in Fig. 1B. More importantly, since the ocular position drift statistics were unchanged across all of our manipulations (Fig. S1, S3C-F), the residual horizontal motions experienced by the neurons would be highly similar to those in the control condition, as if there was no retinal image stabilization at all. Thus, at the individual neural modulation level, horizontal retinal image stabilization should look indistinguishable from full retinal image stabilization relative to the control condition, and vertical retinal image stabilization should appear like the control condition instead; this is despite both conditions causing world-centered grating motions on the display.

Our neurons were modulated according to the predictions of retinotopic pattern displacements over the RF’s (Fig. 1B-E), suggesting sensitivity to local features much smaller than SC RF size. During horizontal retinal image stabilization, the neurons’ activity was elevated as much as in full retinal image stabilization (Fig. 3C, D; average modulation index 20.86%; significantly larger than zero; p=7.8757×10^−9^; 1-sample t-test; n=61 neurons). On the other hand, the neurons’ firing rates were unchanged from control with vertical retinal image stabilization (Fig. 3E, F; population modulation index not significantly different from zero; p=0.3898; 1-sample t-test; n=61 neurons). The difference between horizontal and vertical retinal image stabilization was also robustly evident at the individual neuron level (Fig. S6B), and the spiking statistics in the two conditions mimicked those in either full retinal image stabilization (for the case of horizontal stabilization; Fig. S6C) or control (for the case of vertical stabilization; Fig. S6C). Therefore, SC neurons are sensitive to the image pattern displacements associated with ocular position drifts, despite the small scale of such drifts (and the local image features) relative to RF size. This means that SC neurons with relatively large RF’s can benefit from the edge enhancing properties of ocular position drifts seen perceptually ^13^.

To further demonstrate the sensitivity of SC neurons to minute pattern displacements orthogonal to the local edges in the patterns, we also repeated the same experiments in one monkey but with horizontal, rather than vertical, 4.44 cpd gratings. Thus, in this monkey, we could test some neurons under two different pattern conditions within the same session (Methods). With vertical gratings, we replicated the results of Fig. 3, as seen in Fig. 4A, B. With horizontal gratings (Fig. 4C, D), it was now vertical retinal image stabilization (as well as full stabilization) that resulted in the largest neural modulations; horizontal stabilization (parallel to the now horizontal gratings) caused the least modulations (also see Fig. S7 showing individual neuron firing rates as well as spiking statistics in the different conditions). Thus, it was always ocular position drifts orthogonal to the local pattern orientations that resulted in the largest neural modulations.

**Figure 4.**
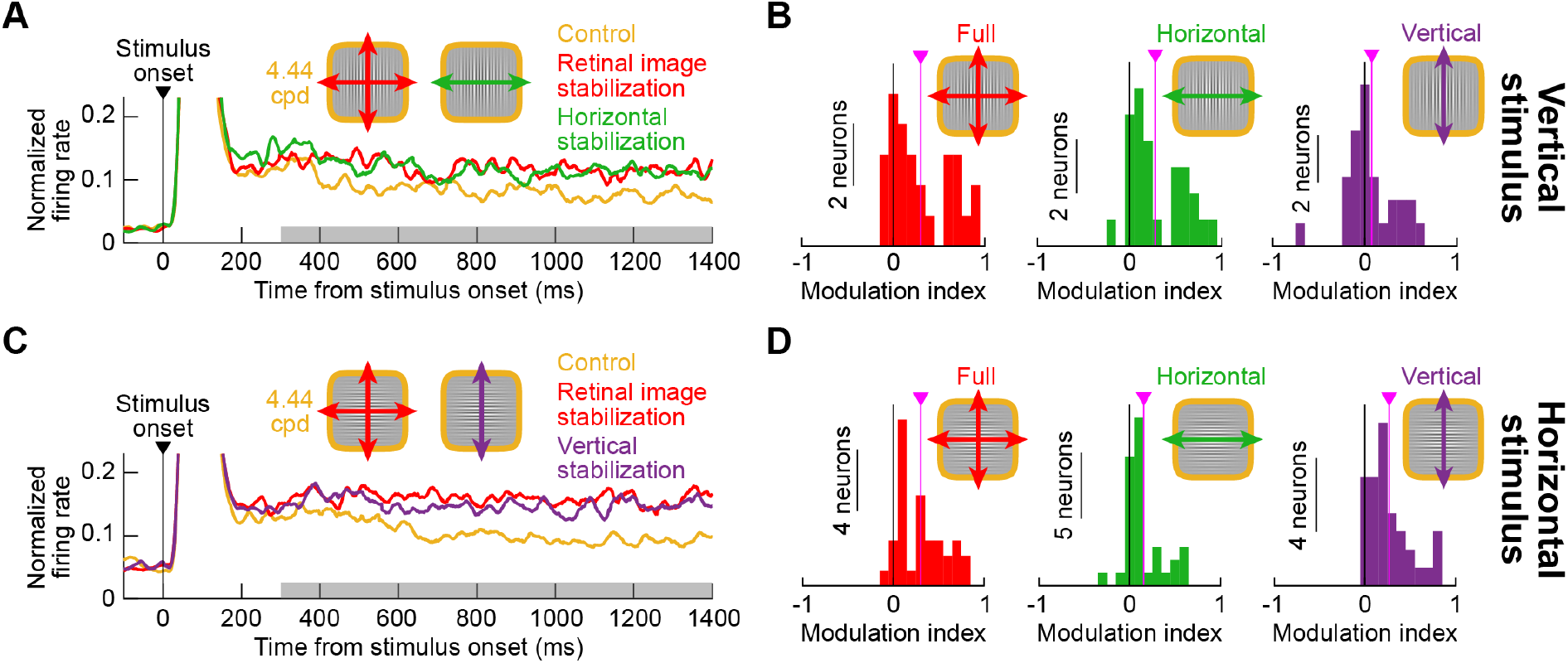
Ocular position drifts orthogonal to a pattern cause the biggest neural modulations, independent of original pattern orientation. **(A, B)** Neural modulations similar to Fig. 3 from a monkey viewing a vertical 4.44 cpd grating. Full and horizontal (orthogonal to the grating orientation) retinal image stabilization caused the biggest neural effects. The modulation indices for vertical retinal image stabilization (rightmost histogram in **B**) were not significantly different from zero (p=0.1728; 1-sample t-test; n=27); the indices were significant for full (p=6.741×10^−5^) and horizontal (p=3.854×10^−5^) stabilization (left and middle histograms in **B**). **(C, D)** SC neural activity modulations in the same monkey viewing a horizontal grating instead of a vertical one. Now, horizontal retinal image stabilization had the weakest effect (middle histogram in **D**; 15.56% modulation; p=1.1×10^−4^; 1-sample t-test; n=35). Vertical retinal image stabilization resulted in similar neural modulations to full retinal image stabilization (26.84% and 30.14%, respectively, in the rightmost and leftmost histograms in **D**; both significantly different from zero: p=1.1×10^−7^ and 4.1×10^−8^). All other conventions similar to Fig. 3. Also see Fig. S7 for individual neuron results.

Therefore, not only are SC neurons sensitive to image pattern displacements on the order of magnitude of 1 min arc, but they are also differentially sensitive as a function of the relative difference between the image displacement directions and the underlying pattern orientations. We previously demonstrated this to be the case in the SC for the image displacements associated with significantly larger microsaccades ^25^, but the smaller scale of ocular position drifts (Fig. 1A, B) suggests an even finer ability of SC neurons to represent and react to detailed visual patterns (Fig. 1B). The SC can indeed contribute to the theoretically-predicted perceptual effects associated with slow ocular position drifts (e.g. ^13,15^).

### Neural responses to high spatial frequency patterns are modulated the most by drift-scale image displacements

The results so far suggest that SC neurons are sensitive to image pattern features that can be significantly smaller than the neurons’ RF sizes (Fig. 1B). However, with a pattern at the optimal RF location, the scale of luminance modulations caused by ocular position drifts (Fig. 1B) should depend on both the spatial detail of the pattern itself as well as the spatial scale of the retinal image displacements caused by eye movements (Fig. S3A, B). Therefore, given the ocular position drift sizes that we observed (Figs. 1A, S1, S3C-F), we expected to observe the largest effects of retinal image stabilization with high spatial frequency patterns (Fig. S3A, B). This was indeed the case. In our experiments, we also tested low (0.56 cpd) and intermediate (2.22 cpd) spatial frequency gratings (Methods). For both vertical (Fig. 5) and horizontal (Fig. S8) gratings, the relevant orthogonal stabilization condition (horizontal in Fig. 5 and vertical in Fig. S8) resulted in the largest neural modulation indices for 4.44 cpd gratings. For example, with vertical gratings and horizontal stabilization, the average modulation index with 4.44 cpd was 20.86% relative to control, but it was 12.46% for 0.56 cpd (p=0.0319; 2-sample t-test; n=61 neurons; comparing 4.44 cpd to 0.56 cpd; Fig. 5B); the modulation index was 13.27% for 2.22 cpd (p=0.0997; 2-sample t-test; n=61 neurons; comparing 4.44 cpd to 2.22 cpd; Fig. 5C). Naturally, full retinal image stabilization also showed similar effects to orthogonal image stabilization, as expected from Figs. 3, 4. Therefore, SC neurons are lawfully sensitive to the luminance modulations caused by orthogonal edges being displaced ever so slightly within their RF’s due to ocular position drifts.

**Figure 5.**
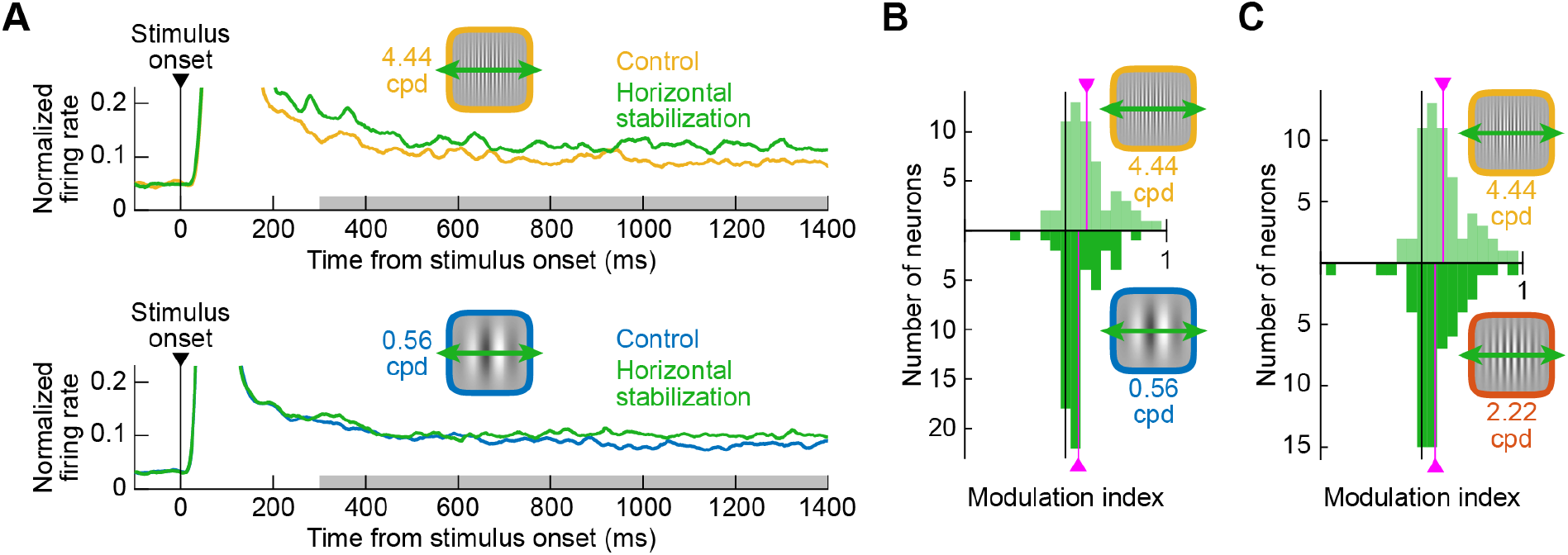
The representation of high spatial frequency features in the SC is most sensitive to the smallest naturally-induced image displacements. **(A)** Normalized population firing rates in control and with horizontal retinal image stabilization for 4.44 cpd (top) and 0.56 cpd (bottom) vertical gratings. The high spatial frequency grating was associated with a larger neural modulation than the low spatial frequency grating (consistent with the image luminance predictions of Fig. S3A, B). **(B)** Neural modulation indices comparing high (top) and low (bottom) spatial frequencies with horizontal retinal image stabilization (pink vertical lines indicate the mean across the population). The high spatial frequency grating was associated with higher modulation indices. **(C)** When comparing the high to the middle spatial frequency, the difference in modulation indices was smaller than in **B** (but the modulation indices with 4.44 cpd were still higher than with 2.22 cpd; 20.86% versus 13.27%).

### Sensitivity to drift-scale pattern displacements still occurs in extrafoveal and lower visual field neurons with larger receptive fields

In prior theoretical and perceptual work ^13,15^, the image pattern consequences of ocular position drifts were primarily, and understandably, considered from a foveal perspective. However, our results above suggest that even eccentricities with larger RF’s (Fig. 1B) may still utilize the visual formatting afforded by slow fixational eye movements in SC visual neural coding. Therefore, we exploited the fact that we sampled neurons from a wide range of eccentricities (Fig. S9), and we specifically analyzed extrafoveal neurons to ask if they were still sensitive to our retinal image stabilization manipulations. These extrafoveal SC neurons clearly showed modulations (Fig. 6A) that were very similar to those observed for the entire neural population (Figs. 3-5): greater elevation with retinal image stabilization for high than low spatial frequencies.

**Figure 6.**
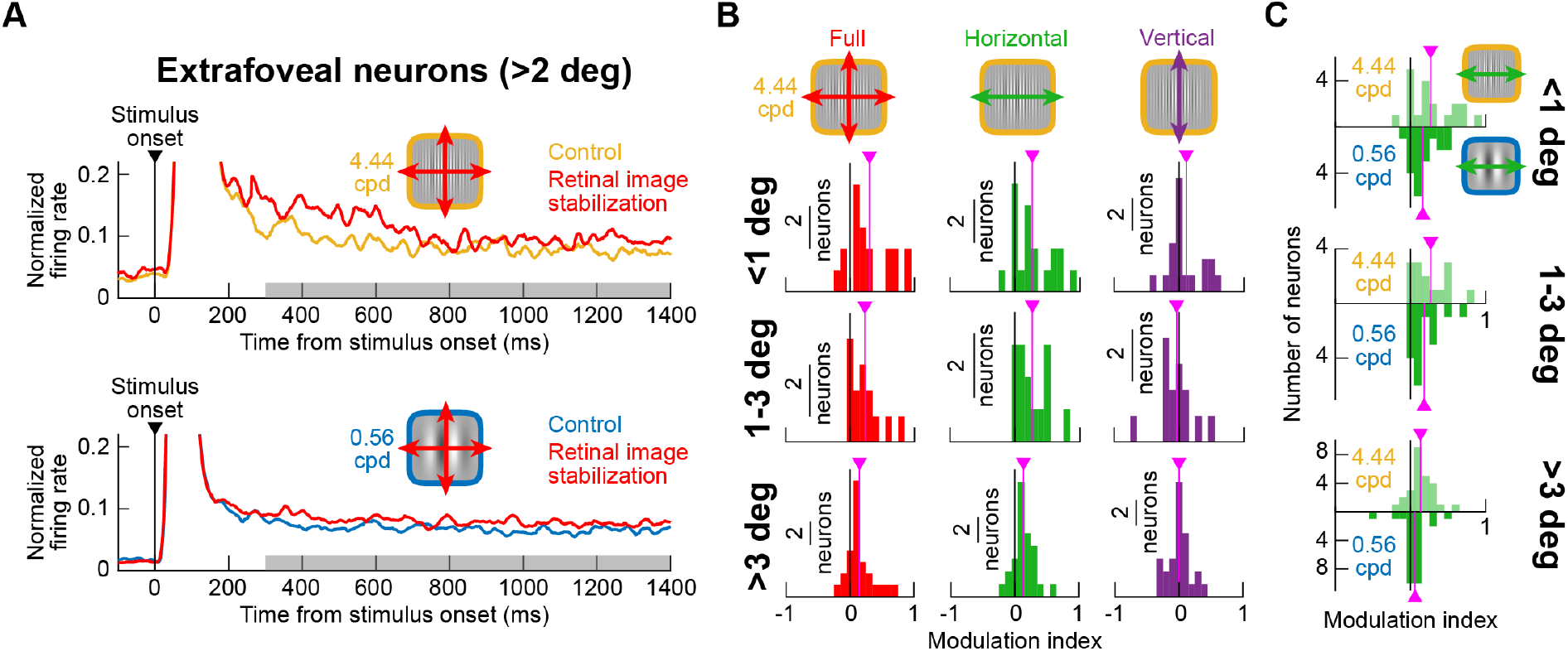
Visual formatting of SC neural responses by ocular position drifts persists for extra-foveal neurons with larger RF’s. **(A)** Normalized population firing rates from all extra-foveal SC neurons comparing control and retinal image stabilization conditions for high (top) and low (bottom) spatial frequencies. Similar observations to Figs. 3-5 could be made: there was still an effect of retinal image stabilization despite the larger RF’s of extra-foveal SC neurons (see also Fig. 1B). **(B)** Modulation indices as in Figs. 3, 4 for three different groups of neurons according to their preferred eccentricities. Full and horizontal retinal image stabilization had positive modulation indices across eccentricities, whereas vertical retinal image stabilization modulation indices were always closest to zero. Note also that the modulation indices for full and horizontal retinal image stabilization progressively decreased in amplitude with increasing eccentricities (top to bottom). **(C)** Modulation indices comparing high and low spatial frequencies (as in Fig. 5) across the different groups of neurons. The same observations as in Fig. 5 were still made for extra-foveal SC neurons.

We then separated the neurons into 3 groups based on their preferred eccentricities (<1 deg, 1-3 deg, and >3 deg). In all cases, full and horizontal retinal image stabilization (with vertical gratings) had higher positive neural modulation indices than vertical retinal image stabilization (Fig. 6B). Specifically, the full and horizontal retinal image stabilization modulation indices in Fig. 6B were statistically significantly different from zero (p<0.0026 in each panel; 1-sample t-test; neuron numbers shown in Fig. 6B), but the vertical retinal image stabilization modulation indices were not. Moreover, the modulation indices were higher for high rather than low spatial frequencies (Fig. 6C); each comparison of high (4.44 cpd) to low (0.56 cpd) spatial frequency modulation indices in Fig. 6C was statistically significant (p<0.032 in each panel; 2-sample t-test; neuron numbers shown in Fig. 6C), except for >3 deg eccentricities (which still showed the same trends; p=0.1683). Thus, even extrafoveal SC neurons with larger RF’s are sensitive to the visual pattern consequences of ocular position drifts on the order of magnitude of 1 min arc, and with the same dependencies as in Figs. 3-5. It should also be noted that the effect sizes of retinal image stabilization were smaller with the larger RF’s (e.g. compare modulation indices across the three rows in Fig. 6B, C), which might be due to the larger integration areas of the larger RF’s in the periphery.

Another test of the effects of RF sizes in our retinal image stabilization manipulations was to also check upper and lower visual field SC neurons. This is so because, in the very same animals, we previously documented a substantial difference in SC RF sizes above and below the horizontal meridian, with lower visual field RF’s being significantly larger ^9^. Here, we found that even such lower visual field neurons, with significantly larger RF’s than upper visual field neurons ^9^, still showed all the same hallmarks of neural modulations described above (Figs. S10, S11). Thus, visual reformatting of the SC neural code by slow ocular position drifts extends well beyond the fovea and affects extrafoveal and lower visual field neurons with larger RF’s (Fig. 1B).

### Neural modulatory effects of ocular position drifts still occur without experimental retinal image stabilization

Finally, if SC neurons are indeed sensitive to the visual pattern consequences of ocular position drifts, might it be possible to observe hallmarks of this even without retinal image stabilization? To test this, we inspected our control trials in more detail. Since we did not have experimental control over individual retinal image stimulation in this condition, we picked, instead, microsaccade-free fixation epochs that were confined within a small eye position window (+/-3 min arc) in any given trial. We hypothesized that, with a vertical grating, momentary epochs of microsaccade-free fixation with particularly large horizontal eye position deviation within such a confined spatial window might drive a luminance transient signal from the neurons more than epochs of low horizontal deviation (a kind of momentary refreshing due to image translation, like with microsaccades ^25^ but on the much smaller scale of ocular position drifts). For example, in Fig. 7A, we had two 250-ms microsaccade-free epochs, with the left one showing a high deviation in horizontal eye position during the first 200 ms and the right one showing a low deviation (Fig. 7B shows the vertical eye position deviations from the same epochs). If we now measured neural activity in the final 50 ms, assuming a temporal integration window of approximately 100-200 ms as per prior measurements of the SC ^26^, then we might expect that the recent history of luminance changes over the RF (in the past 200 ms) provides stronger sensory motion drive from the large horizontal deviation epochs than the low horizontal deviation epochs (or any vertical deviation epochs). In other words, refreshing of the image pattern representation in the RF would be the largest (for a given retinal image position) with an orthogonal recent shift of the pattern.

**Figure 7.**
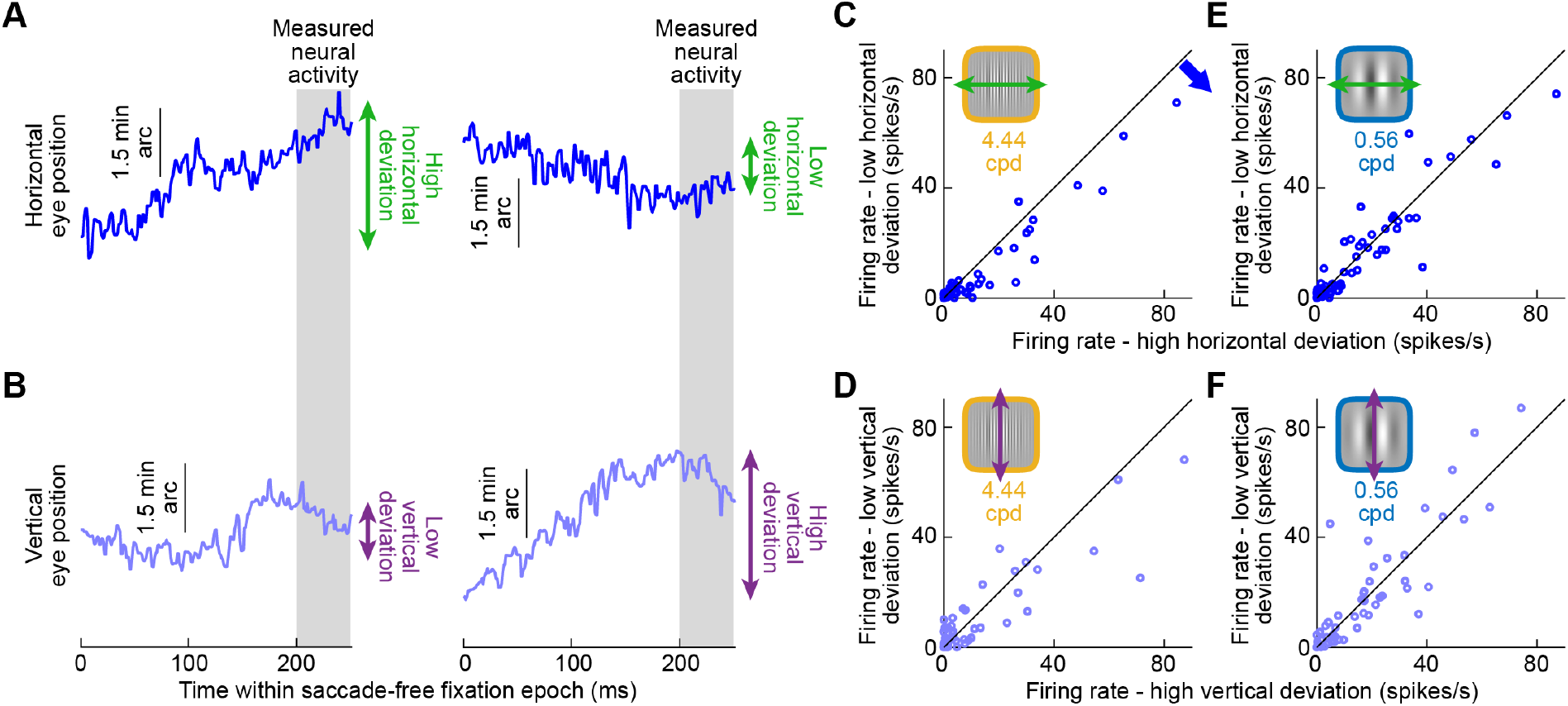
Naturally occurring ocular position drifts in the absence of experimental retinal image stabilization also significantly modulate the representation of high spatial frequency features in the SC. **(A)** Example microsaccade-free fixation epoch from two control trials (right and left) having either a large (left) or small (right) ocular position drift in the horizontal direction. We only picked epochs in which eye position remained within a confined spatial window despite the drift variations (Methods). **(B)** Vertical eye positions from the same two trials as in **A**. The epoch with high horizontal drift had low vertical deviation (left), and the epoch with low horizontal drift had high vertical deviation (right). **(C)** For all control trials with a high spatial frequency grating, we measured firing rate in a 50 ms interval (gray in **A, B**) preceded by a 200-ms microsaccade-free fixation epoch, and we divided the epochs as having a horizontal deviation larger (x-axis) or smaller (y-axis) than the median horizontal deviation across all epochs in a given neuron. Instantaneous firing rates preceded by a period of large horizontal position deviations (orthogonal to the vertical grating) were associated with systematically higher firing rates. **(D)** This effect was absent when assessing the impact of large and small vertical eye position drifts relative to the vertical gratings. **(E, F)** For a low spatial frequency grating, neither horizontal (**E**) nor vertical (**F**) drifts significantly modulated neural activity. For horizontal drifts, this is consistent with the sizes of eye position drifts relative to the gratings’ luminance spatial profiles (Figs. 1, S3A, B). For vertical drifts, this is consistent with the lack of significant luminance modulation of the vertical pattern by a subtle vertical image displacement.

This was indeed the case. We divided all 250-ms microsaccade-free epochs, only from control trials, into high or low horizontal deviation (while still maintaining the average position of the retinal image of the grating within a confined spatial window; Methods), according to the median deviation value in each session. With a vertical grating of 4.44 cpd, the epochs with high horizontal deviation consistently resulted in higher firing rates in the final 50 ms than the epochs with low horizontal deviation (Fig. 7C; p=1.2×10^−4^; 2-sample t-test; n=61 neurons). When we repeated the same analysis based on epochs of large or small vertical eye position deviations instead, there was no longer a difference (Fig. 7D; p=0.1688; 2-sample t-test; n=61). This is because vertical drift over a vertical grating caused minimal image changes in the RF’s, and it is consistent with our horizontal and vertical retinal image stabilization results above (e.g. Figs. 3, 4). Similarly, with low, rather than high, spatial frequency patterns, the effects were significantly diminished (Fig. 7E, F; p=0.445 for E and p=0.8227 for F; 2-sample t-test; n=61 neurons), consistent with Fig. 5. Therefore, SC neurons are sensitive to the visual pattern consequences of ocular position drifts even without any physical stimulus motion on the display caused by our retinal image stabilization technique.

## Discussion

We found that individual SC neurons, even with large RF’s, are sensitive to minute displacements of visual patterns caused by ocular position drifts on the order of 1 min arc in amplitude. We were particularly motivated by the question of whether the diverse visual capabilities of the primate SC, which are becoming increasingly evident, include being sensitive to local feature patterns that are significantly smaller than the RF’s themselves (Fig. 1B). Thus, we investigated whether subtle shifts of these patterns caused by ocular position drifts, representing the lower limit of natural self-induced image displacements, can reliably and systematically modulate SC neural activity. We employed retinal image stabilization to experimentally control the location and motion of a given pattern over the RF at any one moment in time. If the RF’s grossly integrated information all over their spatial extents, then subtle shifts of a given pattern (Fig. 1B) should not have resulted in altered neural responses. In contrast, we found robust neural modulations, which were also direction-dependent. Thus, even though SC neurons may have large RF’s, they still exhibit sensitivity to highly local image features. This idea is consistent with the notion that the primate SC can contribute to visual scene analysis ^4,5,26^ and object processing ^6,27-30^ in a variety of ways.

Our results are relevant with respect to theoretical predictions on how eye position drifts can reformat visual images by introducing temporal fluctuations of image luminance at any one position on the retina ^13-16,31^. Such image reformatting happens by virtue of a physical rotation, albeit small, of the eyeball when viewing a stationary image. However, for such a reformatting to actually influence perception in a meaningful way, then neural elements downstream of the retina must be sensitive to its consequences. With small foveal RF’s, say in the lateral geniculate nucleus or primary visual cortex, this idea might be expected because small ocular position drifts can displace stimuli in and out of the equally small RF’s; that is, the scale of image shifts is similar to the scale of foveal (and perifoveal) RF sizes. However, whether larger RF’s (e.g. extrafoveally), and particularly within an area that is more traditionally investigated from the perspective of motor control like the SC, can still benefit from such reformatting was not equally clear. We found this to be the case, adding to the increasingly rich repertoire of visual capabilities of the primate SC described in the literature. Thus, visual reformatting by ocular position drifts for the benefit of perception can extend also beyond the fovea, and it has direct neural consequences downstream of the retina.

An additional interesting implication of our results concerns the degree of correlation between the two eyes during ocular position drifts. We experimentally stabilized our grating images based on the motion of only one eye (Methods), and we still observed very systematic neural modulations. This means that there must have been at least a minimal amount of correlation between the motions of the two eyes. Otherwise, our retinal image stabilization conditions would have created disparities between the left and right eye images that might have blurred the gratings too much. This would not have necessarily increased the gain and directional sensitivity of neural responses with retinal image stabilization like we saw. Therefore, it will be important in future work to better investigate binocular coordination in ocular position drifts during gaze fixation. Indeed, the question of whether ocular position drifts are correlated (either positively or negatively) between the two eyes has been investigated in the past, with a variety of observations and interpretations ^12,32^. It has also been previously shown that primates are capable of controlling slow eye movements with speeds and position changes similar to those obtained with ocular position drifts during fixation ^20,33,34^.

In yet additional future experiments, stabilization based on only one eye motion could be exploited experimentally to investigate the strength of binocular and monocular visual input integration in the SC. Specifically, it is known that most of the SC is binocular ^1,35-37^. However, it could be that some neurons’ activity may be dominated by one eye input or the other. Therefore, in follow up experiments, one can repeat our study but with stabilization, in separate trials, based on the separate eyes (e.g. stabilization in one trial based on right eye motion and stabilization in another trial based on left eye motion). If a particular SC neuron is functionally dominated by input from one eye, then stabilization with one of the eyes should cause larger modulations than stabilization based on the motion of the other eye. This would allow investigating potential ocular dominance of visual information in individual neurons in the awake monkey SC. Moreover, with sufficient mapping across the SC topographic map using such an experiment, it may then be possible to find individual zones in the SC that are potentially dominated (while remaining generally binocular) by visual input from one eye or the other, similar to the finding of orientation tuning zones in the mouse SC ^38,39^.

Since microsaccades are a component of fixational eye movements, it is also likely that they cause similar visual modulations to ocular drifts, but with different spatial and temporal parameters due to the faster and larger nature of microsaccades. This is exactly what we saw recently in the SC ^25^, and it is also consistent with calculations of the different spatiotemporal consequences of saccades on visual images relative to ocular position drifts ^40^. In any case, it is highly unlikely that microsaccades explain our results in the current study, because we excluded these movements from analysis. Also, the microsaccade characteristics were not altered by our retinal image stabilization manipulations but the neural activity was. Therefore, our experiment isolated the effects of the smaller and slower ocular position drifts on SC activity.

In all, our results highlight the importance of investigating active vision ^41^ in the SC ^42^ and in other visual areas^18,43,44^, whether by careful analysis of the image consequences of eye movements on the retina with stable targets (as in Fig. 7) or by experimentally altering the normal visual-motor loop by gaze-contingent manipulation (as in Fig. 3). This would be even more important by using rich visual stimuli – like gratings, textures, and patterns – and even natural images.

## Acknowledgements

We were funded by the Deutsche Forschungsgemeinschaft (DFG): (1) SFB 1233, Robust Vision: Inference Principles and Neural Mechanisms, TP 11, project number: 276693517; and (2) Werner Reichardt Centre for Integrative Neuroscience excellence cluster (EXC307).

## Author contributions

CYC and ZMH collected the data. ZMH and FK analyzed the data. ZMH wrote the manuscript. ZMH, CYC, and FK edited the manuscript.

## Methods

### Experimental animals and ethics approvals

We recorded superior colliculus (SC) neural activity from two adult, male rhesus macaque monkeys (N and P) aged 7 years, and weighing 8 kg and 7 kg, respectively. The experiments were approved by ethics committees at the regional governmental offices of the city of Tübingen.

### Laboratory setup and animal preparation

The experiments were conducted in the same laboratory as that described in earlier publications ^3,9,45^. Briefly, the monkeys were seated in a darkened booth approximately 45 cm from a calibrated and linearized CRT display spanning approximately +/-22 deg horizontally and +/-15 deg vertically. Data acquisition and stimulus control were managed by a custom-made, real-time computing system ^19,46^, interfacing with the Psychophysics Toolbox ^47-49^ and a Multi-Channel Acquisition Processor (MAP) data acquisition device (Plexon, Inc.).

The monkeys were prepared for behavioral training and electrophysiological recordings earlier ^46,50^. Specifically, each monkey was implanted with a head-holder and scleral search coil in one eye ^46^. The search coil allowed tracking eye movements using the magnetic induction technique ^51,52^, and the head-holder comfortably stabilized head position during the experiments. The monkeys also each had a recording chamber centered on the midline and tilted 38 deg (monkey P) or 35 deg (monkey N) posterior of vertical, allowing access to both the right and left SC ^45^.

### Behavioral task

We employed a gaze fixation task in which we presented static gabor gratings of different spatial frequencies within the receptive fields (RF’s) of the recorded neurons. Unlike in our earlier work with spatial frequency mapping in the SC ^5,53^, we maintained the stimulus on the display for much longer during fixation ^54^. Specifically, once we identified the size and location of an RF, we designed a gabor of suitable size to fill the RF. The grating always had high contrast (100%) and one of three different spatial frequencies (0.56, 2.22, or 4.44 cpd), which were varied across trials (the phase of the gabor was also random from trial to trial).

A trial started with the onset of a central white fixation dot over a gray background. After the monkey fixated the spot in a stable manner for a random interval spanning a few hundred milliseconds, the grating appeared in the RF and remained on for approximately 1500 ms. The fixation spot was small (8.5 × 8.5 min arc), and it had a luminance of 72 cd/m^2^. The gray background had 21 cd/m^2^ luminance. If the monkey successfully fixated the spot for the entire duration of the trial, it was rewarded with fruit juice, and another trial was initiated after a short blank-screen interval.

Four different trial types were interleaved. In the control condition, both the fixation spot and grating were stable on the display. This condition was analyzed for microsaccade-induced visual reafferent responses in a recent study ^25^, as well as for interactions between spiking activity and microsaccade kinematics ^54^. The sustained firing rates during microsaccade-free fixation, as well as all the remaining three trial types of the task, were never described in any other publications. The additional trial types constituted our retinal image stabilization manipulations (Fig. 1). In full retinal image stabilization (Fig. 1C), the grating (from the moment it appeared until trial end) was moved in lockstep with instantaneous eye position (see next section). The fixation spot always remained stable on the display to help anchor gaze properly and not alter the eye movement statistics (Figs. S1-S3). This was necessary because we wanted to isolate the influences of (slow) fixational eye movements on neural activity and, therefore, had to ensure that the eye movements themselves were occurring as naturally as possible. In horizontal retinal image stabilization (Fig. 1D), the vertical position of the grating was kept constant on the display and unchanged from control; the horizontal position of the grating was moved in lockstep with horizontal eye position. Finally, in vertical retinal image stabilization (Fig. 1E), the horizontal position of the grating was stable on the display and similar to the control condition, whereas the vertical position of the grating was moved in synchrony with vertical eye position.

In a subset of experiments, we replaced the vertical gratings with horizontal ones, in order to explore the relative relationship between eye movement directions and image pattern orientations (e.g. Fig. 4).

We collected approximately 18 trial repetitions per condition per neuron.

### Retinal image stabilization

We first calibrated eye position measurements using methods described earlier ^19^. Briefly, at the beginning of every session, the monkeys fixated (multiple times) a series of 19 locations on the display for at least 1000 ms. We then measured raw voltages during stable fixation from each location. To convert the raw voltages to degrees of angular rotation, we used a multi-order polynomial including both the horizontal and vertical raw voltages, as well as cross-channel interaction terms ^19^.

We then employed our real-time gaze-contingent display system ^19,20,46^. In this system, we sampled and processed eye positions at 1 KHz using a real-time control system, and we updated the display at 120 Hz (constrained by the display technology). In retinal image stabilization trials, after every display refresh time, we sampled new eye positions and processed them. We then calculated the position of the grating according to the new eye positions, and we updated the display at the next frame refresh. Thus, our retinal image stabilization trials discretized eye positions at 120 Hz (the bottleneck imposed by the display refresh rate). Such a rate is suitable for successful retinal image stabilization with slow eye movements, as shown previously by our ^19,20^ and other ^13,17,55,56^ laboratories. In fact, even with microsaccades, Fig. S5 shows that the microsaccade-related reafferent responses were reduced by retinal image stabilization at 120 Hz. This means that while the microsaccades were still a bit too rapid for the display’s 120 Hz frequency, the stimulus was still moved sufficiently rapidly to catch up with the real microsaccade and reduce the retinal slip of the grating by the eye movement (and, therefore, the associated reafferent response).

Finally, eye coil systems often exhibit a slow drift in their measurements, which is much slower than ocular position drifts (we confirmed this by comparing initial fixation positions across trials and assessing a time constant of eye coil system drift, which was more than two orders of magnitude slower than within-trial ocular position drifts). Therefore, at the beginning of every trial, we performed a drift correction that was applied for all subsequent eye position measurements within a trial. We did this by averaging eye position in the final 50 ms before gabor onset and using this measurement as a reference to which we compared all subsequent eye positions in the trial. In post-hoc analyses, if there was a microsaccade in the drift correction measurement interval, the trial was excluded from further analysis.

### Eye movement data analysis

We detected saccades and microsaccades as described previously ^25,46,57^. We used the detections for two primary purposes. First, we established that microsaccade properties were not altered by our retinal image stabilization manipulations (Fig. S2). Second, for our neural analyses, we excluded all data starting from 10 ms before microsaccade onset until 90 ms after microsaccade end. This was done in order to avoid movement-induced reafferent responses from the analyses (Fig. S5). We also inspected all trials for blinks, and we removed all blink intervals, including a period of 50 ms before and a period of 50 ms after each of them.

Our microsaccade analyses (Fig. S2) included microsaccade rate, directions, and main sequence ^58^ relationship between movement amplitude and peak velocity. For microsaccade rate, we used a running window of 25 ms width and moved in steps of 2 ms. Within each such time window, we counted the fraction of trials in which the window contained microsaccades across trials of a given condition. For microsaccade directions and main sequence relationships, we considered all microsaccades in the sustained fixation interval that we were interested in for our neural analyses (>300 ms after stimulus onset; see *Neural data analysis* below), and we plotted the distribution of movement vector angles (for the direction analysis) or the scatter of movement peak velocity versus movement radial amplitude (for the main sequence analysis). We performed all of these analyses separately for each of the four stimulus conditions (control and three retinal image stabilization versions), and we then compared the results to confirm that retinal image stabilization did not significantly alter the statistics of microsaccades.

To confirm that retinal image stabilization also did not alter absolute eye position (i.e. the combination of ocular position drifts and microsaccades), we also plotted the raw eye positions (in sustained fixation; >300 ms after stimulus onset) across all trials in each condition (Fig. S1).

We also analyzed ocular position drifts more specifically. For example, for Fig. S3C-F, we first excluded all microsaccades and their pre- and post-movement periods mentioned above. We then considered all sustained fixation intervals starting from 300 ms after stimulus onset in each trial. For every interval in between two successive microsaccades (which we called a saccade-free fixation interval), we measured mean eye position and subtracted it from every sample of eye position within the same interval. This gave us the instantaneous deviation of eye position from the mean position during the particular saccade-free interval of interest. In Fig. S3C, E, we then binned all such deviations across all fixation intervals and obtained a distribution of how much ocular position drifts altered eye position. For Fig. S3D, F, we also took the standard deviation of eye position within each microsaccade-free fixation interval. We then plotted the distribution of these measurements across all such intervals in Fig. S3D, F. Our purpose in both cases was to highlight that the slow ocular position drifts had similar characteristics whether we ran control or retinal image stabilization trials (Fig. S3C-F), and also to predict the amounts of luminance modulations over the retina that were expected from our gratings and ocular position drift amplitudes (Fig. S3A, B).

### Neural data analysis

We recorded from 61 individually isolated SC neurons, which were first characterized online using delayed visually-guided and memory-guided saccade tasks ^9,45,53^. The initial characterization allowed classifying the neurons as being visual or visual-motor, as well as assessing the neurons’ RF sizes and locations. After establishing that a recorded neuron was visually-responsive, we ran the main behavioral task described above with the grating placed at the optimal RF position (except for some test examples like in Fig. S4). We included all visually-responsive neurons in our analyses, without further classification into visual or visual-motor categories. This was because our results were similar regardless of whether a neuron was purely visual or visual-motor in nature (the results were also highly consistent across the population; for example, Fig. S6A, B).

In both monkeys, we tested the neurons with vertical gratings. In monkey N, we additionally tested 35 neurons with horizontal gratings (27 of these neurons had both vertical and horizontal gratings tested together within the same session).

We analyzed neural data by counting spikes during the sustained fixation interval or by converting spike times into firing rate estimates (with a Gaussian convolution kernel of σ 40 ms). We defined the sustained fixation interval as the time from 300 ms to 1400 ms after grating onset. The lower bound of this time interval (300 ms) was chosen to avoid the initial visual onset response of the neurons (occurring immediately after grating onset); the upper bound was chosen to maximize the numbers of neurons that we could pool in the analyses. We did not notice a difference in the onset response strength (the initial visual burst after grating onset) across our different retinal image stabilization manipulations relative to control. Therefore, we did not analyze initial visual responses further. Rather, we were interested in assessing how subtle image displacements (i.e. during sustained presence of a stimulus within the RF’s) affected SC neural activity. Our chosen interval of more than 1 second (300 ms to 1400 ms from stimulus onset) was sufficient to do that.

As stated above, in all of our analyses, except for Fig. S5, we excluded all neural activity associated with microsaccades, in order to avoid contamination by microsaccade-induced reafferent responses ^25^ (Fig. S5 shows an example of such responses). We replaced all intervals starting from 10 ms before microsaccade onset to 90 ms after microsaccade end by not-a-number labels such that these intervals were not included when computing across-trial averages of firing rates or when computing inter-spike intervals. Note that all of our neurons were not microsaccade-related in the sense of emitting a motor burst at movement onset; they, therefore, did not exhibit prolonged buildup of discharge up to 100 ms before microsaccade onset ^59,60^. This justified our choice of pre-microsaccadic mask interval (also see Fig. S5A).

For summarizing population firing rates (e.g. Fig. 3A), we first calculated the within-neuron average firing rate across trial repetitions of a given spatial frequency in the control condition. We then normalized each trial’s firing rate (for the same spatial frequency) by dividing the trial’s instantaneous firing rate (at any given time after stimulus onset) by the peak of the average firing rate in the first 150 ms after stimulus onset. That is, we normalized each trial’s firing rate to the peak average firing rate occurring in the early stimulus-evoked visual burst interval of the control condition. We repeated this trial-by-trial normalization procedure for all control trials of the same spatial frequency, and also for all trials of the retinal image stabilization conditions (again of the same spatial frequency). Thus, if a retinal image stabilization condition (e.g. full retinal image stabilization) elevated firing rates relative to control, then the normalized firing rates were also elevated. With all trials and all spatial frequencies normalized to each neuron’s response in the respective control condition, we could then average normalized trials across neurons to get population firing rates like in Fig. 3A. This allowed us to visualize the neural modulation effects of retinal image stabilization, and to do further individual neuron and population analyses.

Among such analyses, we calculated neural modulation indices (e.g. Fig. 3B). To do so, we measured raw firing rates at the individual neuron level across conditions. For each trial of a given spatial frequency, we evaluated the average sustained firing rate (300 ms to 1400 ms after stimulus onset, and with microsaccades removed as per the procedure described above). We did this in the control condition, and we averaged all measurements across trial repetitions (*fr*_*control*_). We then repeated this procedure for one of our retinal image stabilization manipulations (full, horizontal, or vertical retinal image stabilization) to obtain *fr*_*stabilization*_ (the average within-neuron sustained firing rate during retinal image stabilization). Then, we calculated a modulation index comparing retinal image stabilization to control as: (*fr*_*stabilization*_ - *fr*_*control*_)/(*fr*_*stabilization*_ + *fr*_*control*_); that is, the modulation index was the average sustained firing rate in a retinal image stabilization manipulation minus the average sustained firing rate in the control condition, divided by the sum of the two measurements.

We plotted histograms of modulation indices across the entire population of neurons, with indications of the average modulation index across neurons (e.g. pink vertical line in Fig. 3B). We tested whether the population modulation indices were significantly different from zero using a t-test and reported p-values in the figures and/or text. In the Results text, we also often reported the average modulation index values as percentages by multiplying the values from the calculation above by 100. We also often compared neural modulation indices across conditions (e.g. comparing the effects of retinal image stabilization for high or low spatial frequency gratings). We did so by performing a t-test across the different conditions and reporting p-values.

For scatter plots of individual neuron results, we followed similar procedures to those above for the modulation indices. For example, in Fig. S6A, we counted the number of spikes per trial in the sustained interval (i.e. per 1100 ms occurring between 300 ms and 1400 ms after stimulus onset) in either the control condition or the full retinal image stabilization condition. This resulted in a paired measurement per neuron (average spike count per control trial and average spike count per retinal image stabilization trial). We then plotted all of these measurements across the population as a scatter plot. Note that microsaccades were still excluded in such analyses, exactly as above. However, since microsaccade characteristics were unchanged across conditions (Fig. S2), the microsaccade exclusion was not inappropriately favoring one condition over the other when comparing the spike counts across them. Therefore, it was appropriate to exclude the occasionally occurring rapid eye movements in this manner. Also note that the spike counts in these kinds of analyses were essentially estimates of average sustained firing rates. This is so because such spike counts were evaluated over an interval of approximately 1 second duration (1100 ms).

We also computed inter-spike intervals during sustained fixation in either control or one of the retinal image stabilization manipulations (e.g. Fig. S6C). Because we excluded microsaccades, there could be intervals within trials that were replaced with not-a-number labels during the exclusion process. Therefore, for inter-spike interval measurements, we first found contiguous blocks of fixation data within trials that existed in between successive microsaccades. Then, we computed inter-spike intervals within all such contiguous blocks. This allowed us to avoid counting inter-spike intervals during (and around) microsaccades, and also to avoid having erroneously large inter-spike intervals due to the not-a-number labels introduced during preprocessing (e.g. if we had counted two spikes on either side of a not-a-number block of data).

For summarizing microsaccade-induced reafferent responses at the individual neuron level in Fig. S5B, we used a similar procedure to our earlier analyses (e.g. Fig. S6A). The only difference is that our measurement interval was now different. Here, for each microsaccade in a given condition, we measured the firing rate in the interval 30-80 ms after microsaccade onset (see gray bar on the x-axis in Fig. S5A). This interval captured the reafferent response. We then compared, within each neuron, the average response with and without retinal image stabilization (Fig. S5B).

For some of our analyses, we selected neurons according to their preferred eccentricities or visual field locations. For example, we categorized neurons based on whether they were foveal or extrafoveal (Fig. 6); or whether they were part of the upper or lower visual field representation of the SC (Figs. S10, S11). To do so, we classified neurons according to the eccentricity and direction from horizontal of their RF hotspots (Fig. S9).

Finally, for Fig. 7, our goal was to ask whether horizontal or vertical ocular position drifts in control trials were still sufficient to modulate SC neural responses in a manner that was consistent with the retinal image stabilization results. For every control trial of a given spatial frequency, we had a moving window of 250 ms duration (starting from 300 ms after stimulus onset and in steps of 1 ms). If the window was devoid of microsaccades (including the pre- and post-microsaccadic masks described above) and the eye position was within +/-3 min arc (horizontally and vertically) from the initial fixation position, we measured eye position variability (horizontal or vertical) in the first 200 ms of the interval and firing rate in the final 50 ms of the interval. That is, we assumed that SC neurons integrate the recent history (200 ms) of the image over the RF’s in their instantaneous firing rate. This was justified based on prior measurements of the temporal properties of SC neurons ^26^. Our measure of variability was the standard deviation of eye position during the interval. We then did a median split based on this variability across all microsaccade-free epochs of a given neuron, and we compared firing rates for high or low variability trials. We did this independently for horizontal and vertical variability. Thus, for vertical gratings, we could compare epochs with high variability of horizontal eye position (orthogonal to the grating) to epochs with low variability of horizontal eye position. If our retinal image stabilization results were indeed related to the image luminance modulations of orthogonal eye position shifts, then such comparison would yield a difference between high and low horizontal drift variability. Similarly, if we now compared epochs of low and high vertical eye position variability, we would have expected no neural effects (analogous to parallel retinal image stabilization). We did this procedure for all neurons and across different spatial frequencies.

## Supplementary figures

**Figure S1.**
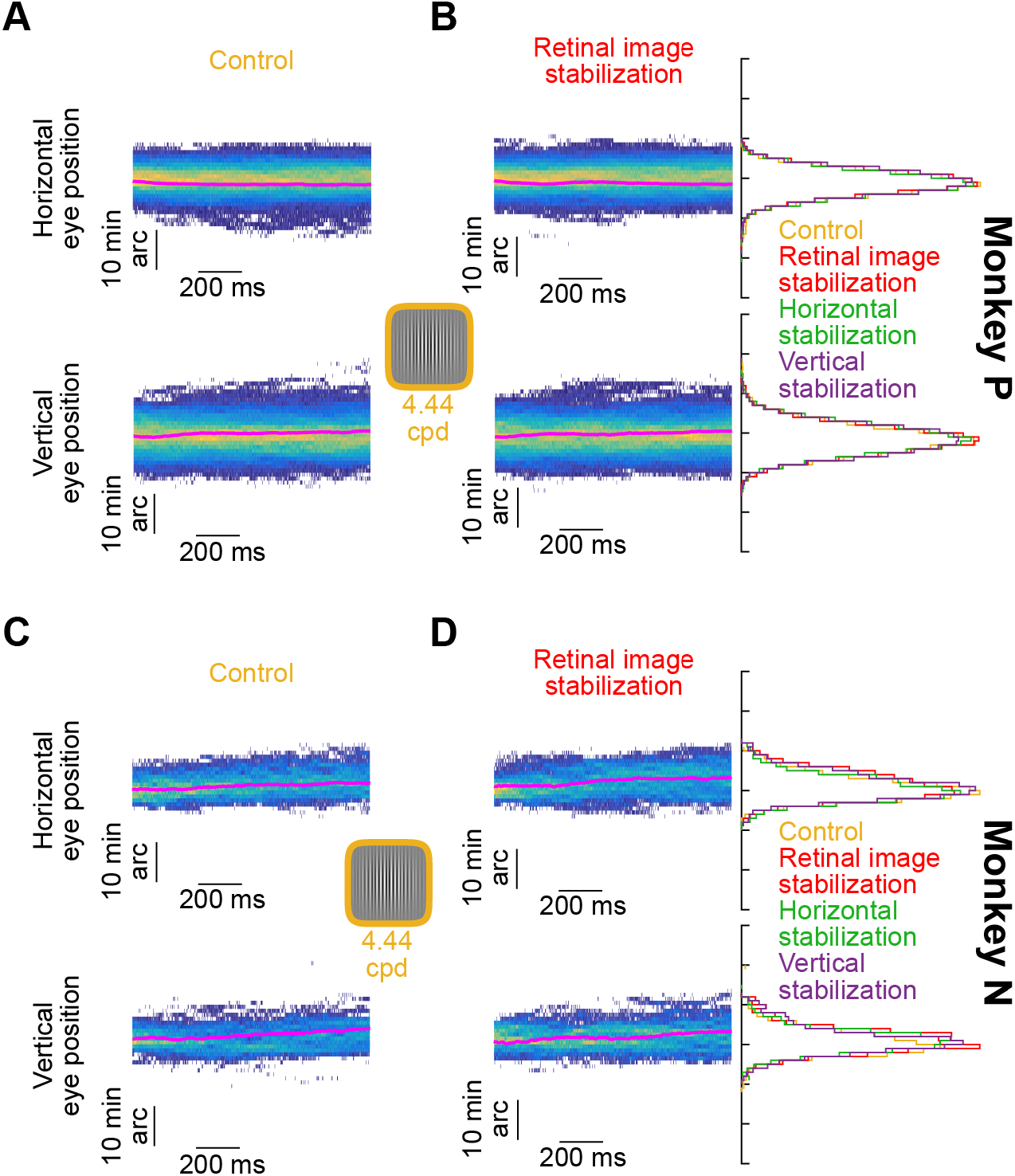
Similarity of fixational eye positions during retinal image stabilization and control trials. **(A)** Histograms of raw horizontal (top) and vertical (bottom) eye positions from monkey P as a function of time from stimulus onset, in the interval from 300 ms to 1400 ms (control trials with a 4.44 cpd gabor grating are shown). The thick pink line in each histogram is the mean eye position. **(B)** Same as **A** but for full retinal image stabilization. The marginal histograms on the right show the distributions of eye positions (across all times) for the control condition, the retinal image stabilization condition (red), as well as the two other retinal image stabilization conditions of Fig. 1D, E. The distributions of eye positions were similar across all conditions, suggesting that retinal image stabilization did not alter eye movement characteristics in our experiments (Methods). **(C, D)** Same as **A, B** but for monkey N. The same results were observed in both monkeys for the low (0.56 cpd) and intermediate (2.22 cpd) spatial frequency trials.

**Figure S2.**
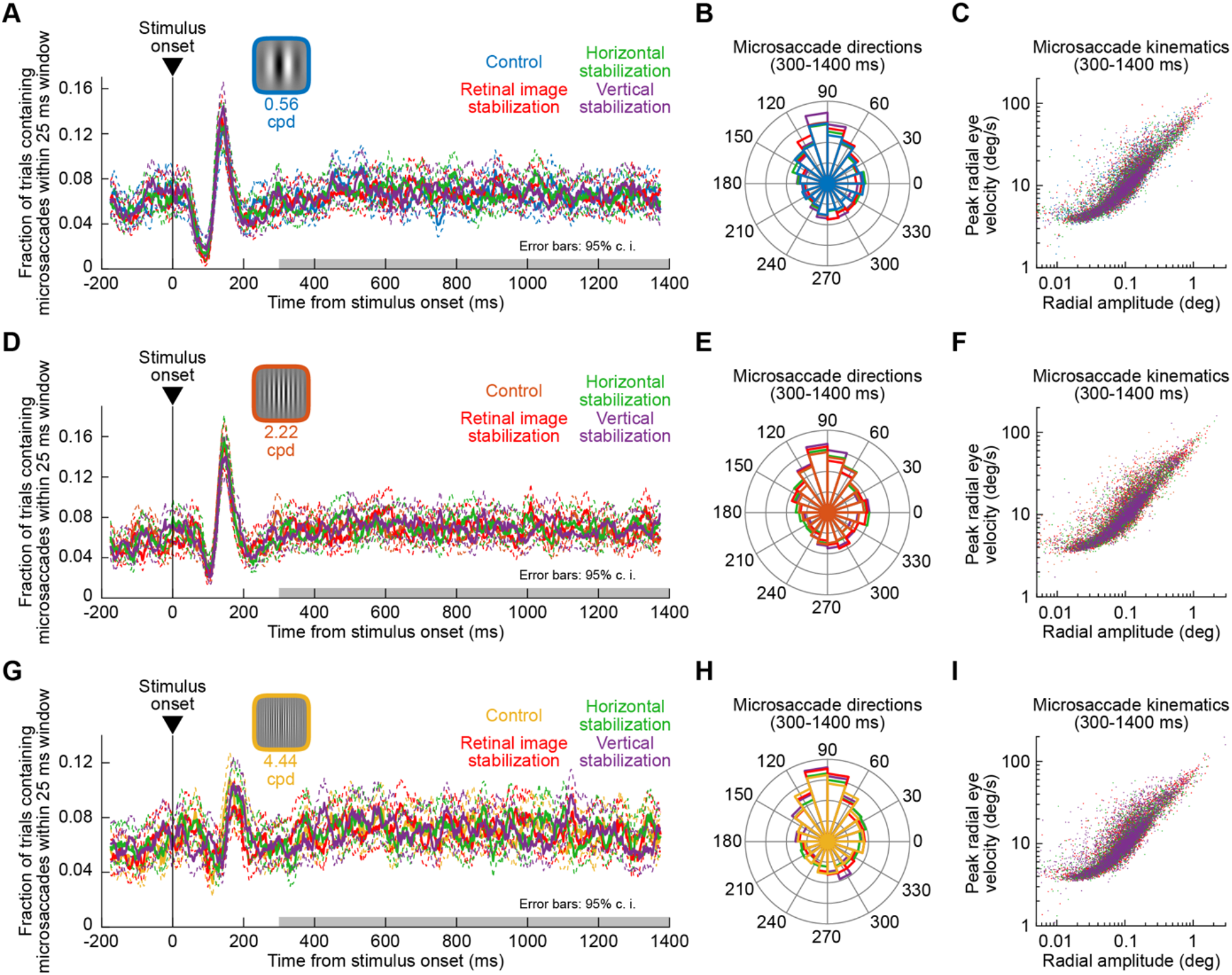
Similarity of microsaccade characteristics between control trials and all retinal image stabilization trials. **(A)** Microsaccade rate (+/-95% confidence intervals) as a function of time from stimulus onset from all trials with a 0.56 cpd grating (monkey P shown as an example). The different colors show the different conditions (control plus the three retinal image stabilization conditions of Fig. 1C-E). Initially, microsaccade rate was modulated by stimulus onset, as expected ^50,61,62^, and was then stable. Critically, the modulations in microsaccades were similar across all of our experimental conditions (control versus the three types of retinal image stabilization). The gray bar denotes our analysis interval for exploring the influences of microsaccade-free ocular position drifts on SC neural activity. **(B, C)** Microsaccade directions (**B**) and kinematics (**C**) during sustained fixation were also unaltered by retinal image stabilization. **(D-F)** Same as **A**-**C**, but for the trials in which a 2.22 cpd grating was presented. **(G-I)** Same as **A**-**C**, but for 4.44 cpd trials. Monkey N showed the same results of no impact of retinal image stabilization on microsaccade characteristics. Note that in all of our neural analyses (except for Fig. S5), we excluded all microsaccades as well as pre- and post-movement intervals around them before taking any measurements (Methods). Our purpose in the current analysis was merely to demonstrate that the properties of microsaccades (like drifts in Fig. S1) were largely unaffected by the retinal image stabilization technique. This was due to the stability of the fixation spot on the display.

**Figure S3.**
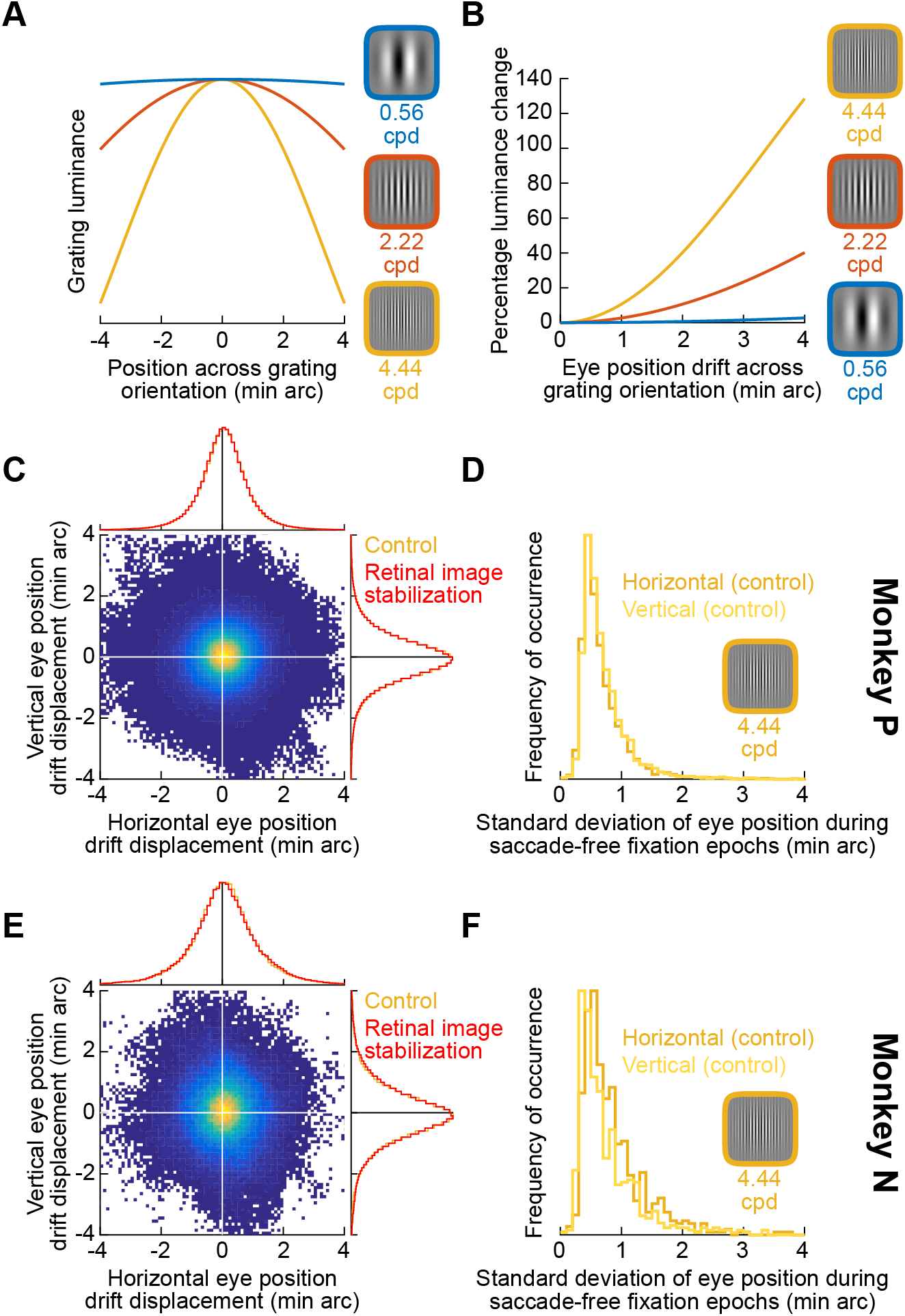
Spatial scale of the image pattern displacements over SC neurons’ visual RF’s. **(A)** Luminance profile of each of our three tested spatial frequencies as a function of position orthogonal to the grating orientation. **(B)** Ocular position drifts on the scale of 1 min arc are associated with the highest instantaneous change in luminance experienced at a given retinotopic location for the highest spatial frequency grating. **(C)** Two dimensional histogram of naturally-occurring displacements of eye position during microsaccade-free fixation. Within any microsaccade-free epoch, we measured the range of eye position deviation from mean position during the epoch (Methods). We did this for the control condition from one of our trial types (4.44 cpd trials), but the results were the same across all conditions (see Figs. S1, S2). The horizontal/vertical yellow histograms show the marginal distributions in each direction; the red histograms overlaid on top show the same distributions from the full retinal image stabilization condition for comparison. Microsaccade-free fixation epochs were associated with displacements on the order of 1 min arc, and were unchanged by our gaze-contingent manipulation (compare red and yellow histograms). **(D)** Variability estimate of microsaccade-free eye position drifts in monkey P (retinal image stabilization conditions yielded similar histograms). **(E, F)** Similar observations in monkey N. Natural fixation behavior in both monkeys was expected to cause predictable luminance modulations of image patterns over SC receptive fields, and such behavior was unchanged by our experimental manipulations (see Figs. S1, S2).

**Figure S4.**
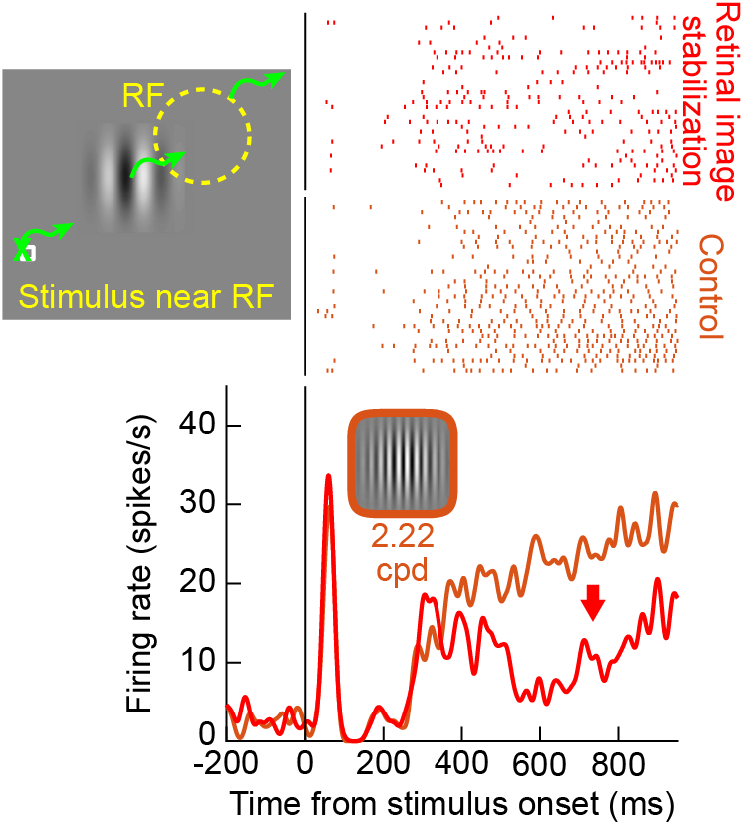
Validating the retinal image stabilization technique by forcing a grating position at a sub-optimal RF position (away from RF center). The schematic shows the experimental manipulation that we applied for this neuron. We presented a grating stimulus displaced away from the preferred RF location indicated by the dashed yellow circle. Therefore, the stimulus was near the RF, but at a sub-optimal position. Forcing this position during retinal image stabilization significantly reduced the sustained response of the neuron. This is the complement of the example neuron results shown in Fig. 2, in which forcing a stimulus at the best receptive field location elevated the neural response relative to control.

**Figure S5.**
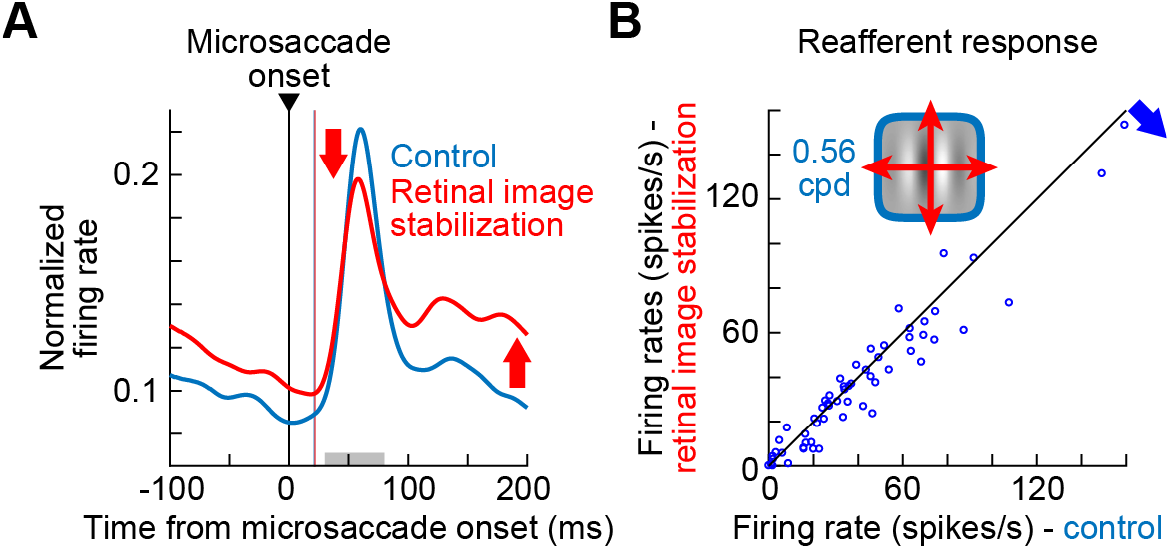
Validating the retinal image stabilization technique by exploring SC visual reafferent responses after microsaccades ^25^. **(A)** Across all neurons, we plotted normalized firing rate (as in Fig. 3A), but this time by aligning data to microsaccades during sustained fixation (as opposed to during microsaccade-free fixation, as in Fig. 3). We did this for presentations of a 0.56 cpd vertical grating. In control trials, there was an expected visual reafferent response immediately after microsaccades ^25^. With retinal image stabilization, firing rate was elevated long before and long after microsaccades (consistent with our main results like in Figs. 2, 3). However, the reafferent response was reduced relative to control. This is because, even though microsaccades were fast relative to display updates, the retinal image stabilization technique still partially tracked these rapid eye movements. This resulted in subdued retinal-image motion caused by the microsaccades (relative to the control condition). Such a reduction in microsaccade-induced retinal-image motion is known to reduce the SC visual reafferent response ^25^. **(B)** Across all neurons, we measured the microsaccade-induced reafferent response (during the gray interval on the x-axis in **A**) in the control and retinal image stabilization trials (Methods). There was a consistent reduction in the reafferent response during retinal image stabilization (p=0.0015; 2-sample t-test; n=61 neurons). Higher spatial frequency gratings had significantly weaker reafferent responses even during control trials ^25^, making the effects of retinal image stabilization harder to observe.

**Figure S6.**
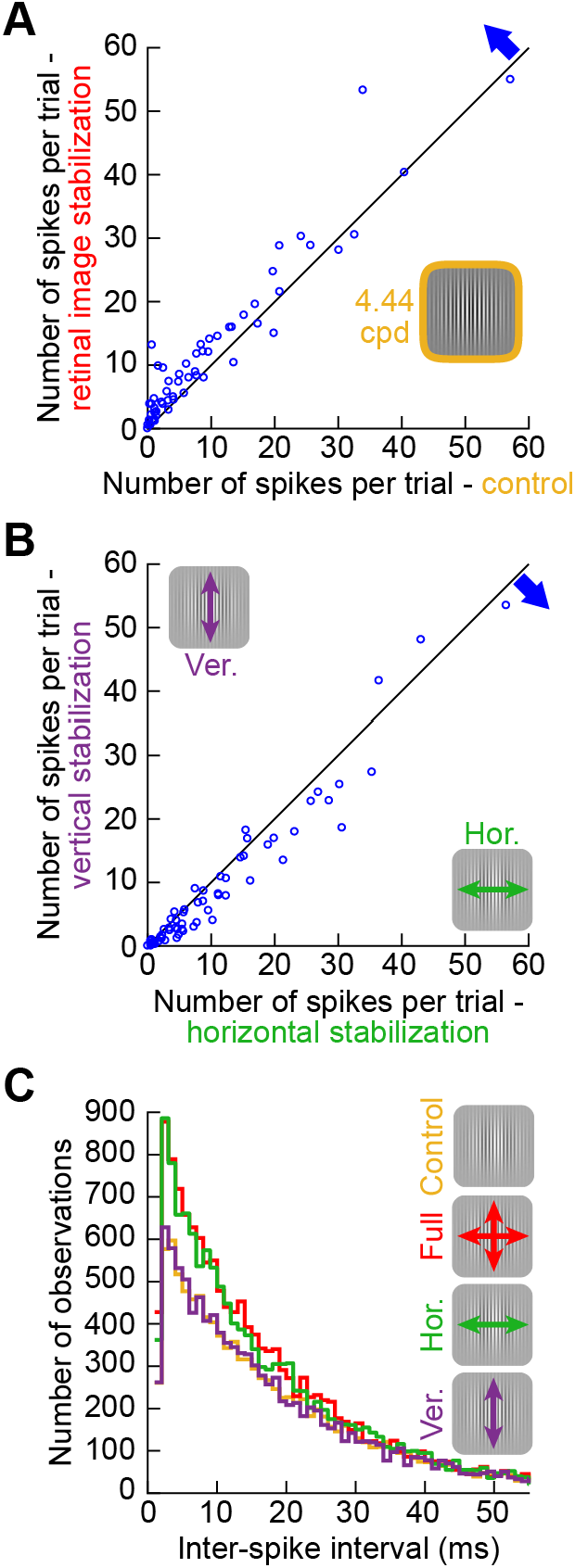
Individual neuron results from Fig. 3. **(A)** We measured raw neural activity during sustained fixation (excluding peri-microsaccadic intervals; Methods) for the analyses of Fig. 3A, B. The x-axis counts the number of spikes per trial in control, and the y-axis counts the number of spikes per trial in retinal image stabilization. Note that given the length of our measurement interval (gray region on the x-axis in Fig. 3A), the shown values of spike counts are quantitatively equivalent to the approximate sustained firing rate of the neurons during the trials. Also, note that since microsaccade characteristics were unaltered by retinal image stabilization (Fig. S2), the shown differences in firing rates across the conditions could not be attributed to potentially different distributions of microsaccades across image stabilization manipulations. Rather, there was a consistent elevation of sustained neural activity (blue arrow) by retinal image stabilization (p=1.436×10^−5^; 2-sample t-test; n=61 neurons). This is consistent with the results of Fig. 3A, B. **(B)** We also compared the individual neuron spike counts across the horizontal and vertical retinal image stabilization conditions (as in Fig. 3C-F). The neurons consistently exhibited elevated activity for horizontal (orthogonal) rather than vertical retinal image stabilization (p=3.142×10^−6^; 2-sample t-test; n=61 neurons). **(C)** Across all neurons, inter-spike interval distributions reflected the results of Fig. 3: horizontal and full retinal image stabilization resulted in more SC activity than in control trials, but vertical retinal image stabilization did not. The inter-spike interval distributions also reveal that the SC spiking statistics (e.g. burstiness) were not significantly altered by retinal image stabilization.

**Figure S7.**
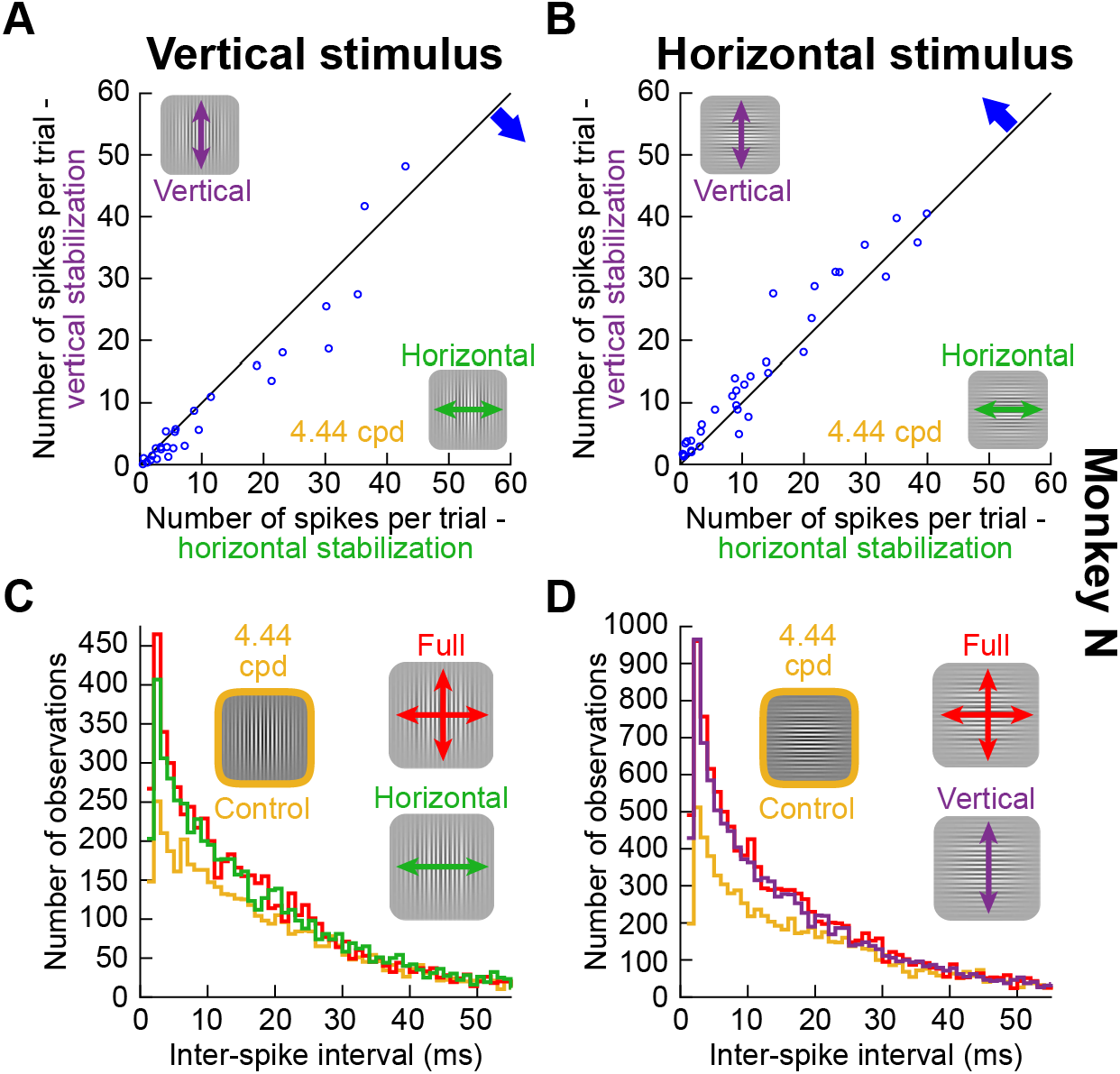
Individual neuron results with horizontal gratings showed similar results to the main experiments with vertical gratings. **(A)** Same as Fig. S6B but from the monkey in which we also collected horizontal grating trials. The same results as in Fig. S6B were observed, as expected (p=0.0072). **(B)** In the same animal, when we flipped the image pattern from vertical to horizontal, it was now vertical retinal image stabilization trials that resulted in elevated firing rates relative to horizontal retinal image stabilization trials (p=0.0026; 2-sample t-test; n=35 neurons). Therefore, it was the relative (orthogonal) relationship between the image pattern and the stabilization direction that mattered for the neurons, consistent with Figs. 4, S8. **(C, D)** Similar to Fig. S6C but for vertical (**C**) or horizontal (**D**) gratings in the same animal. Note that with horizontal gratings (**D**), it was now vertical retinal image stabilization that resulted in indistinguishable results from full retinal image stabilization (instead of horizontal retinal image stabilization as in the case of vertical gratings in **C**).

**Figure S8.**
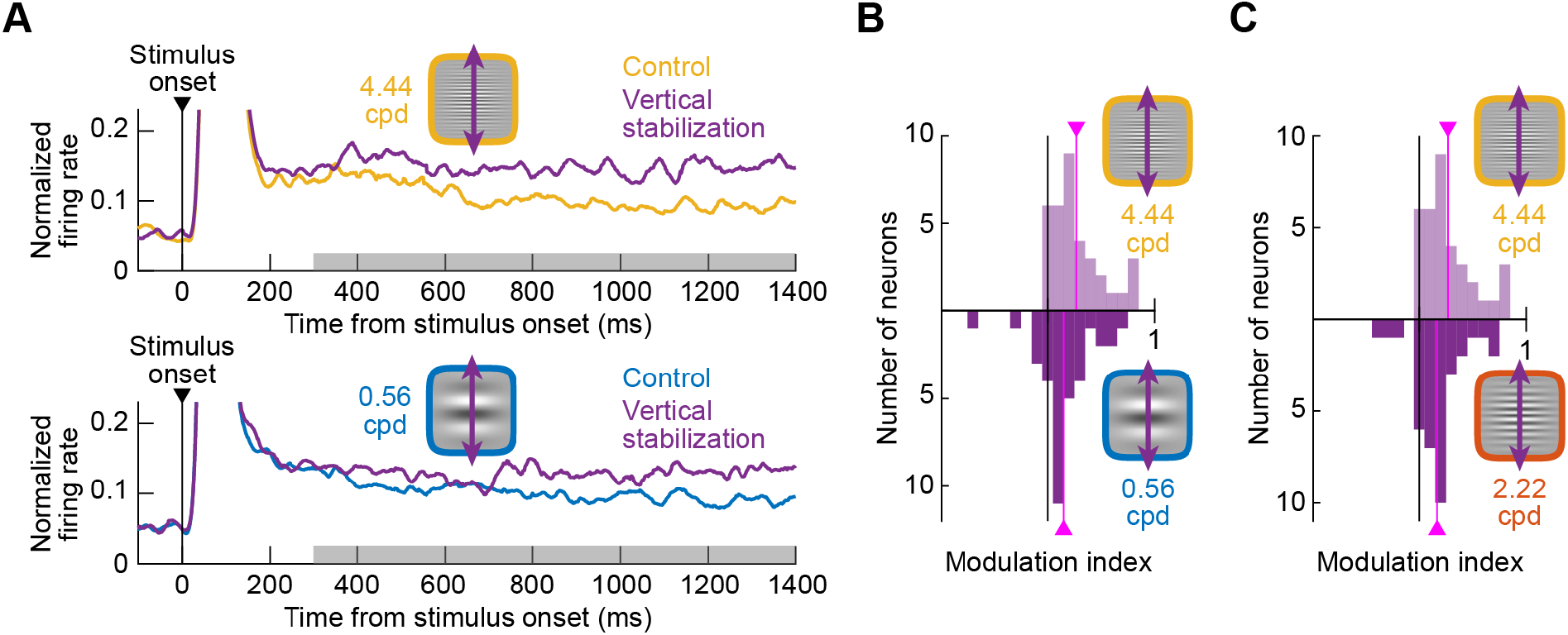
Dependence of neural modulation on spatial frequency even with horizontal gratings. **(A)** Similar to Fig. 5A but for horizontal gratings and vertical retinal image stabilization (Fig. 4 showed that vertical retinal image stabilization with horizontal gratings was equivalent to horizontal retinal image stabilization with vertical gratings due to the orthogonal relationship between eye movements and image patterns in both cases). There was a higher modulation of neural activity by the retinal image stabilization technique for high (top) than low (bottom) spatial frequencies (compare each stabilization curve to its respective control curve), consistent with Fig. 5. **(B)** Similar to Fig. 5B but for horizontal gratings with vertical retinal image stabilization. The population average modulation indices were 26.84% and 14.94% for 4.44 cpd and 0.56 cpd gratings, respectively (p=0.0529; 2-sample t-test; n=35 neurons). **(C)** Similar to Fig. 5C but for horizontal gratings and vertical retinal image stabilization. Similar results to **B** were now obtained when comparing 4.44 cpd (26.84% average modulation index) to 2.22 cpd (16.76% average modulation index; p=0.0845 comparing 4.44 cpd to 2.22 cpd modulations; 2-sample t-test; n=35 neurons). Therefore, with both vertical (Fig. 5) and horizontal (this figure) gratings, the same dependence of neural activity on the relative spatial scale of image patterns and ocular position drifts was observed.

**Figure S9.**
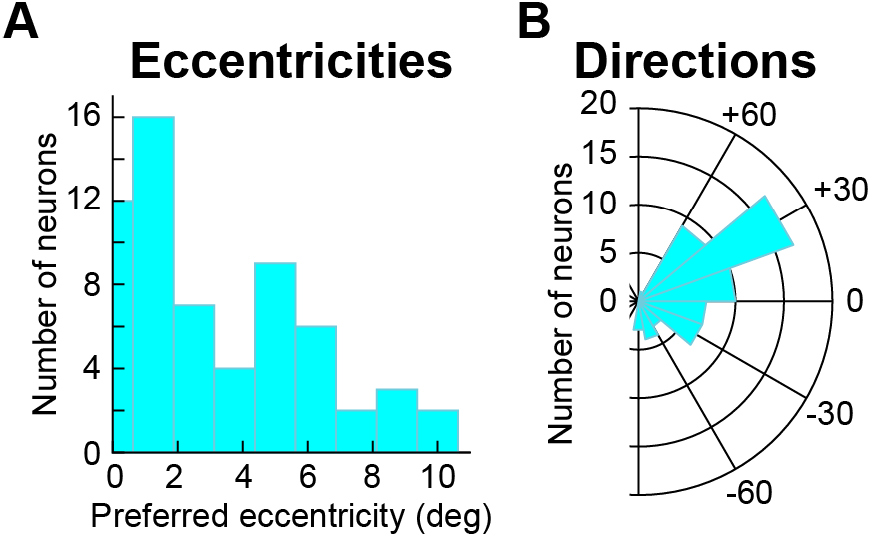
RF hotspot locations of the recorded neurons. **(A)** Distribution of preferred eccentricities by our neurons. We sampled foveal and extrafoveal neurons. **(B)** Distribution of the directions of the RF hotspot locations relative to the horizontal meridian. We sampled both upper and lower visual field neurons.

**Figure S10.**
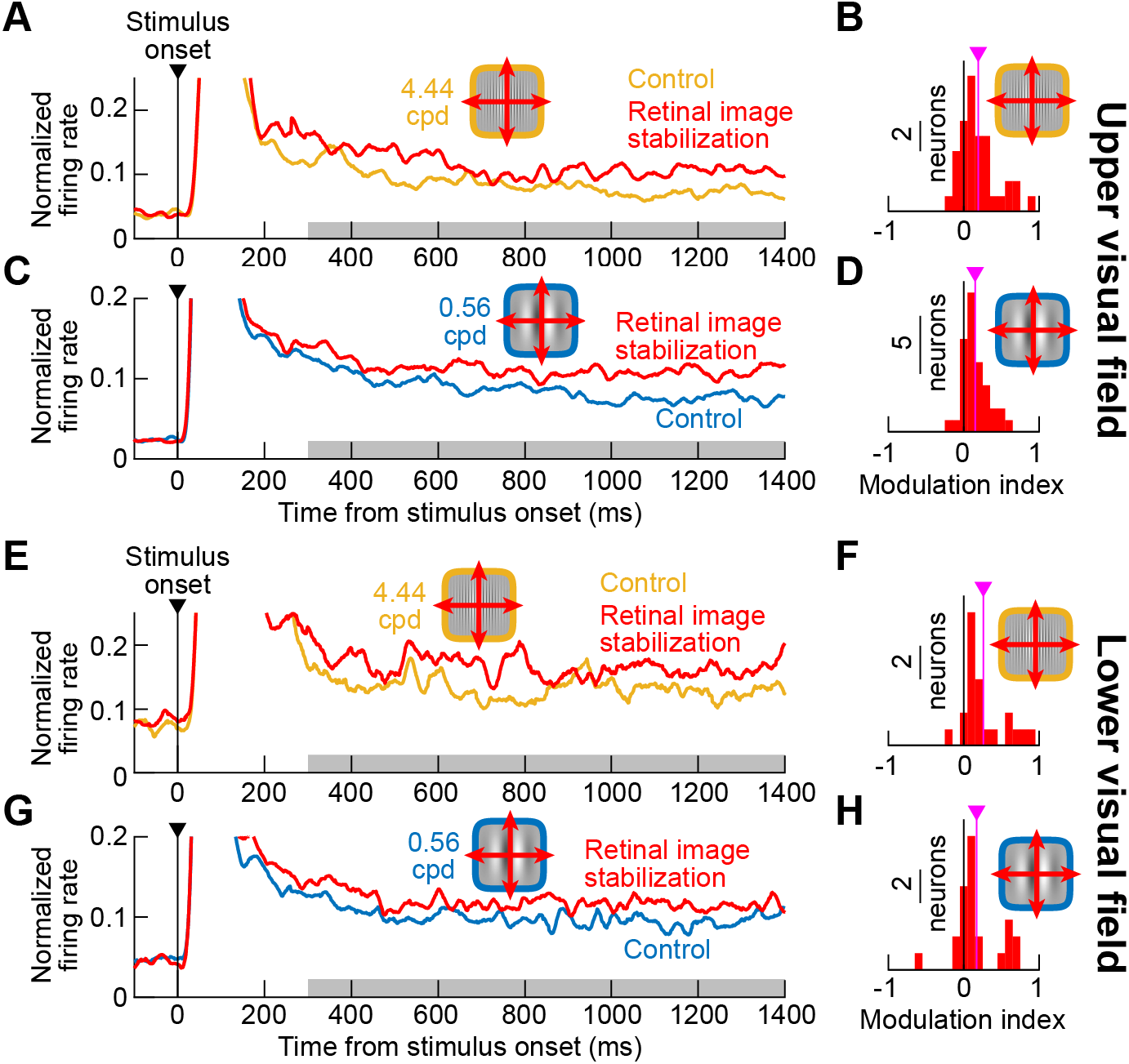
Sensitivity of both upper and lower visual field SC neurons to the visual-pattern consequences of ocular position drifts on the scale of 1 min arc amplitude. **(A)** Normalized firing rates in control and retinal image stabilization from all neurons with RF hotspots occupying the upper visual field. A 4.44 cpd vertical grating was presented to the neurons. As in Fig. 3, neural activity was systematically elevated with retinal image stabilization. **(B)** Modulation indices for the individual neurons in **A**. There was a significant positive modulation across the population (average modulation of 19.2% across the population; pink vertical line; p=4.009×10^−5^; 1-sample t-test; n=37 neurons). **(C, D)** Same as **A, B** but for a low spatial frequency grating. The average modulation index was now 15.13% (p=1.649×10^−6^; 1-sample t-test; n=37 neurons). **(E-H)** Same as **A**-**D** but for lower visual field SC neurons, which we showed earlier (in the same animals) to have significantly larger RF’s than upper visual field neurons ^9^. There was still significant positive elevation of neural activity for these neurons (p=2.94×10^−4^ for **F** and p=0.0134 for **H**). Therefore, even SC neurons with relatively large RF’s are still sensitive to the visual-pattern consequences of ocular position drifts.

**Figure S11.**
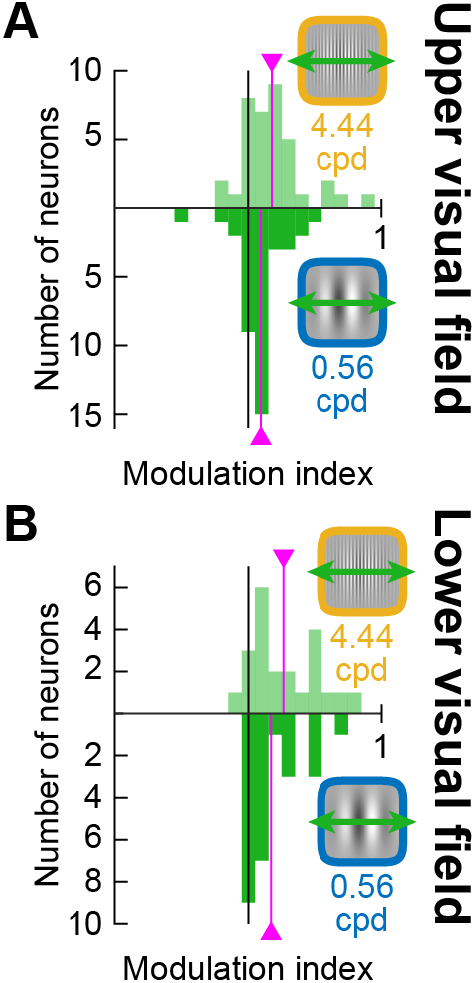
Ocular position drifts cause stronger modulations with high spatial frequency patterns than low spatial frequency patterns in both upper and lower visual field SC neurons. **(A)** Similar analyses to Fig. 5 but only for upper visual field neurons. The average modulation indices (pink vertical lines) for high and low spatial frequencies were 17.61% and 9.49%, respectively (p=0.0481; 2-sample t-test comparing high to low populations; n=37 neurons). **(B)** Same as **A** but for lower visual field neurons with larger RF’s. The modulation indices were now 26.33% and 17.03% for high and low spatial frequencies, respectively (p=0.0066; 2-sample t-test comparing high to low spatial frequency populations; n=24 neurons). Similar results were obtained with full retinal image stabilization, as expected.

